# Geographic variation and bias in polygenic scores of complex diseases and traits in Finland

**DOI:** 10.1101/485441

**Authors:** Sini Kerminen, Alicia R. Martin, Jukka Koskela, Sanni E. Ruotsalainen, Aki S. Havulinna, Ida Surakka, Aarno Palotie, Markus Perola, Veikko Salomaa, Mark J. Daly, Samuli Ripatti, Matti Pirinen

**Affiliations:** Institute for Molecular Medicine Finland, Helsinki Institute of Life Sciences, University of Helsinki, Finland; Analytic and Translational Genetics Unit, Massachusetts General Hospital, Boston, USA; Stanley Center for Psychiatric Research, Broad Institute of MIT and Harvard, Cambridge, USA; Program in Medical and Population Genetics, Broad Institute, Cambridge, USA; National Institute of Health and Welfare, Helsinki, Finland; Department of Internal Medicine, University of Michigan, Ann Arbor, MI, US; Psychiatric & Neurodevelopmental Genetics Unit, Department of Psychiatry, Massachusetts General Hospital, Boston, USA; Department of Neurology, Massachusetts General Hospital, Boston, USA; Department of Public Health, University of Helsinki, Helsinki, Finland; Helsinki Institute for Information Technology HIIT and Department of Mathematics and Statistics, University of Helsinki, Helsinki, Finland

## Abstract

Polygenic scores (PS) are becoming a useful tool to identify individuals with high genetic risk for complex diseases and several projects are currently testing their utility for translational applications. It is also tempting to use PS to assess whether genetic variation can explain a part of the geographic distribution of a phenotype. However, it is not well known how population genetic properties of the training and target samples affect the geographic distribution of PS. Here, we evaluate geographic differences, and related biases, of PS in Finland with geographically well-defined sample of 2,376 individuals from the National FINRISK study. First, we detect geographic differences in PS for coronary artery disease (CAD), rheumatoid arthritis, schizophrenia, waits-hip ratio (WHR), body-mass index (BMI) and height, but not for Crohn’s disease or ulcerative colitis. Second, we use height as a model trait to thoroughly assess the possible population genetic biases in PS and apply similar approaches to the other phenotypes. Most importantly, we detect suspiciously large accumulation of geographic differences for CAD, WHR, BMI and height, suggesting bias arising from population genetic structure rather than from a direct genotype-phenotype association. This work demonstrates how sensitive the geographic patterns of current PS are for small biases even within relatively homogenous populations and provides simple tools to identify such biases. A thorough understanding of the effects of population genetic structure on PS is essential for translational applications of PS.

## Introduction

Understanding the causes behind geographic health differences can help to optimally target the limited health care resources and improve public health. Geographic health differences can be partially explained by life style and environmental factors but also by genetic differences that affect health both through population specific genetic diseases, e.g. the Finnish disease heritage (The Finnish Disease Heritage), and through variation in the polygenic components of many complex diseases (Fuchsberger et al. 2016; Hartiala et al. 2017; O’Connell et al. 2018). In particular, recent discoveries from genome-wide association studies (GWAS) (Visscher et al. 2017) have enabled improved polygenic prediction of complex diseases and traits and raised expectations for their future translation to clinical use (Mavaddat et al. 2015; Abraham et al. 2016; Khera et al. 2018; Stocker et al. 2018). An open question is to which extent the geographic distribution of phenotypes could be explained by their polygenic predictions.

A standard way to estimate a polygenic score (PS) of an individual is to select a set of independent variants identified by a GWAS, to weight the number of copies of each variant by its effect size estimate from the GWAS, and to sum these quantities over the variants. PS have turned out to be a useful tool for identifying high risk individuals in many diseases such as breast cancer (Mavaddat et al. 2015), prostate cancer (Schumacher et al. 2018) and Alzheimer’s disease (Stocker et al. 2018). As an example, a PS for coronary artery disease (CAD) can characterize individuals with risk equivalent to carrying a monogenic variant of familial hypercholesterolemia (Khera et al. 2018). At the same time, two recent studies have raised concerns about comparing PS between populations with varying demographic histories (Martin et al. 2017; Reisberg et al. 2017). Both studies showed that when a PS was built on a GWAS conducted in European populations and then applied to populations from Africa or East Asia, the differences in the PS were inconsistent with the actual phenotypic differences between the populations. Exact reasons for this inconsistency are unclear, but it has been speculated that a complex interplay of population genetic differences, including varying linkage-disequilibrium patterns and allele frequency differences, between the target sample and the GWAS data can limit generalizability across populations (Martin et al. 2017; Reisberg et al. 2017). Can similar problems appear also within a much more genetically and environmentally homogeneous setting than between populations from different continents? This is a crucial question for the public health care systems in countries that have the growing potential to implement polygenic scores as part of their population-wide practice.

In this work, we evaluate the geographic distribution of PS of several complex diseases and traits in Finland and demonstrate how the effect of genetic population structure needs to be assessed before PS can become a robust tool for population-wide use. The data resources available in Finland provide several favorable characteristics for this study. First, on a world-wide scale, Finland has a demographically and socially homogeneous population and a top-level public health care system (GBD 2016 Healthcare Access Quality Collaborators 2018), which together reduce many possible environmental effects contributing to geographic variation in health. Second, some notable geographic differences in phenotypes and general health still occur in Finland. A good example is the incidence rate of CAD that is 1.6 times higher in eastern Finland than in western Finland (THL) (Figure 1). In fact, even larger differences in CAD incidence were observed in the 1970s and despite the extensive and successful public health campaign to reduce these rates through the Northern Karelia project (Puska et al. 2009), differences between east and west still remain today. Third, the genetic structure in Finland is well-characterized (Lappalainen et al. 2006; Jakkula et al. 2008; Salmela et al. 2008; Neuvonen et al. 2015; Kerminen et al. 2017; Martin et al. 2018b) (Figure 1), which enables a detailed comparison between the geographic distribution of PS and the overall genetic population structure within the country.

**Figure 1.**
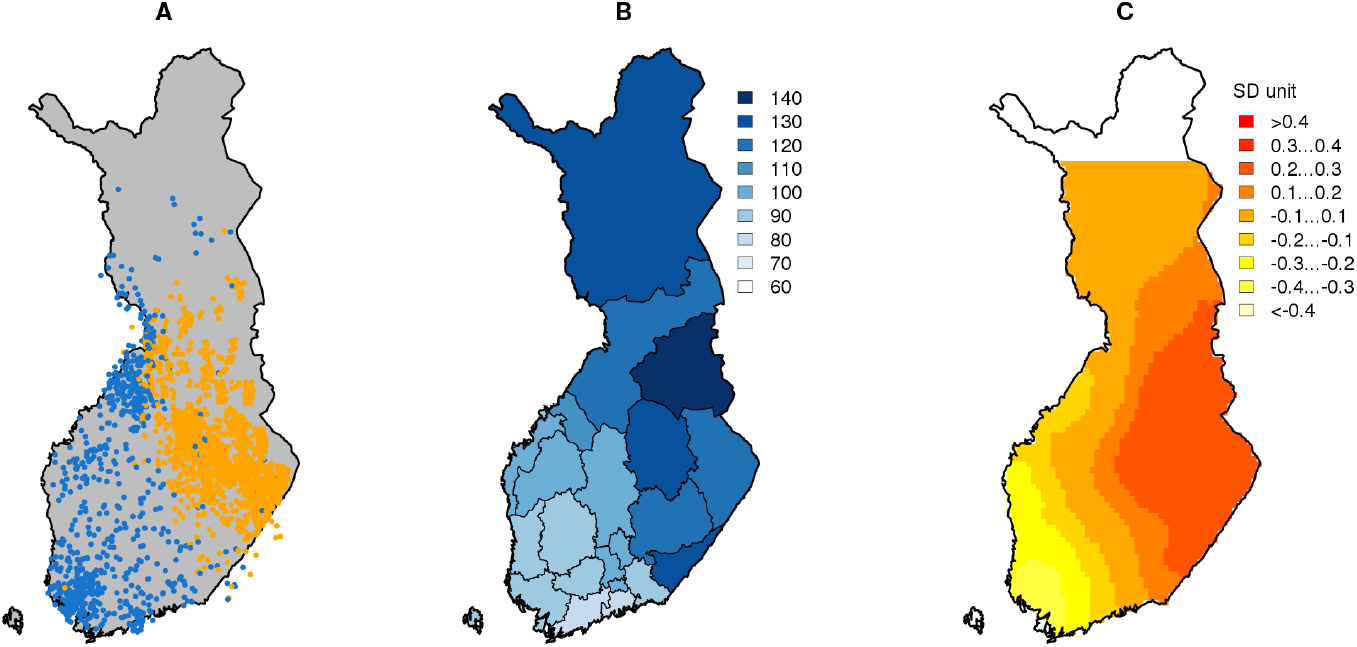
**A)** Main genetic population structure, **B)** incidence rate for age adjusted coronary artery disease (CAD), 2013-2015 (THL, Sepelvaltimotauti-indeksi) **C)** distribution of polygenic score (PS) for CAD in Finland. Population structure was estimated by clustering 2,376 samples into two groups (Kerminen et al. 2017). Incidence rate is scaled to have a mean = 100. PS distribution is shown in unit of standard deviation.

In our analyses, we observe clear geographic structure in PS distributions for most phenotypes considered. Furthermore, the spatial pattern is similar across the phenotypes and resembles the population genetic east-west division of Finland (see comparison for CAD in Figure 1). While a population genetic difference can well result in such patterns, a major goal of this work was to thoroughly assess whether these geographic patterns could alternatively result from some bias emerging when the GWAS estimates of tens of thousands of variants are accumulated into PS. We do this by generating many versions of PS with different inclusion criteria of variants and by monitoring how the geographic structure accumulates across these PS.

To demonstrate our approach, we consider the adult height (HG) as a model trait. In addition to HG, we apply our approach to two additional quantitative traits: body-mass index (BMI) and waist-hip ratio (WHR) and five diseases: coronary artery disease (CAD), rheumatoid arthritis (RA), schizophrenia (SCZ), Crohn’s disease (CD) and ulcerative colitis (UC). The results suggest that polygenic components of CAD, RA, SCZ, WHR, BMI and HG show differences along the east-west direction, while only HG and WHR also show differences in the north-south direction. PS for CD and UC do not show significant regional differences in either direction. Last, we discuss the credibility of the observed geographic differences. In particular, we report possible population stratification-related biases in PS for CAD, WHR, BMI and HG. Our results raise concerns about how to reliably interpret geographic variation in PS even within relatively homogeneous populations.

## Results

### Polygenic scores show geographic differences in Finland

We estimated PS across Finland using a geographically well-defined sample of 2,376 individuals from the National FINRISK Study 1997 survey (Borodulin et al. 2017). Each of these 2,376 individuals had his/her parents born within 80 km from each other and the mean of the parents’ coordinates were used as the individual’s location. We derived PS for the individuals using summary statistics from publicly available GWAS meta-analyses by applying linkage disequilibrium pruning (r^2^ < 0.1), minor allele frequency filtering (MAF > 0.01) and P-value thresholding (P < 0.05) (Methods). To visualize the results on the map of Finland (Figure 2), we estimated the score at each map point by averaging individuals’ PS inversely weighted by the individuals’ squared distance from the point (Methods).

**Figure 2.**
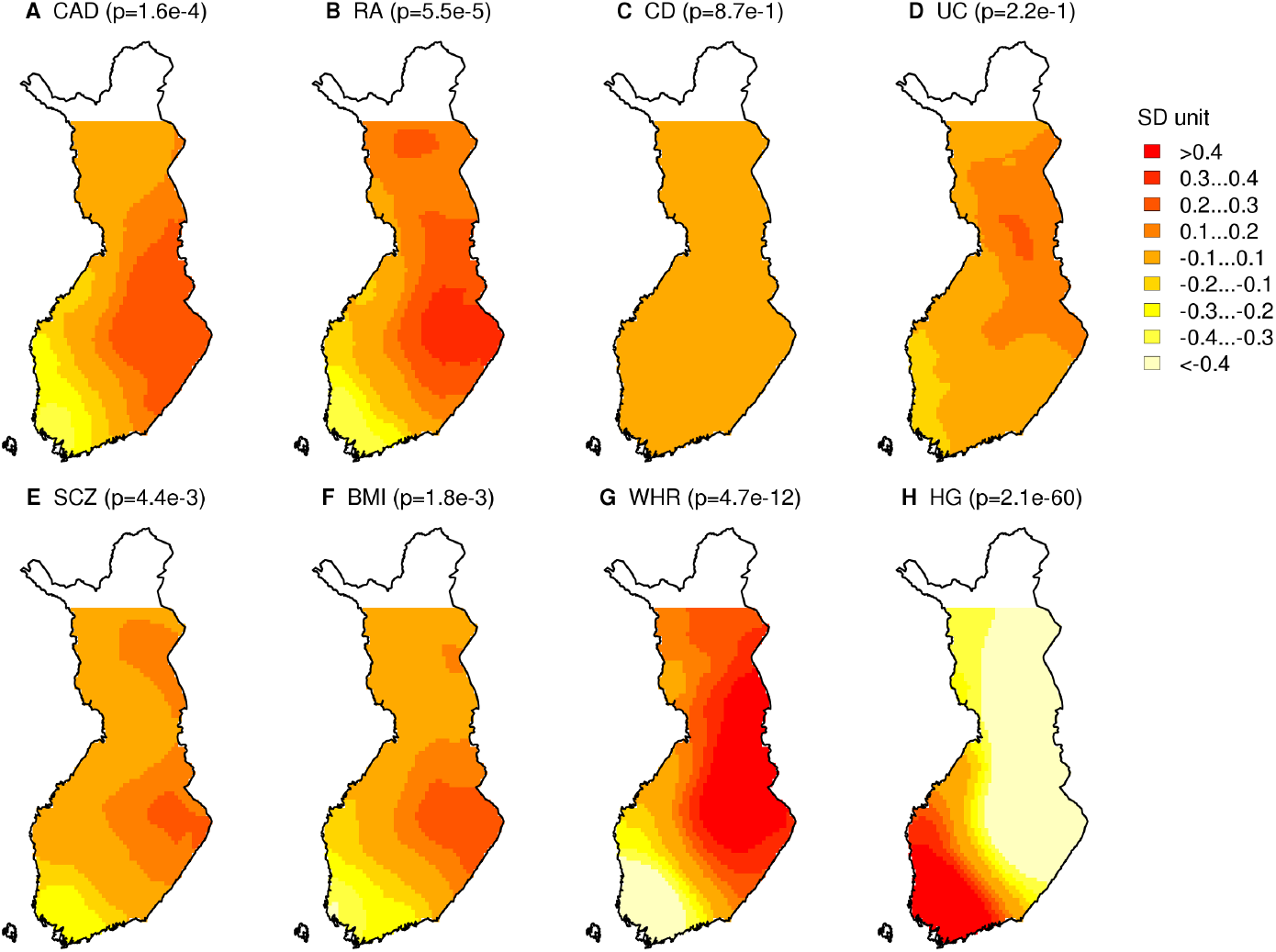
Distribution of polygenic scores for **A)** Coronary Artery Disease, **B)** Rheumatoid Arthritis **C)** Crohn’s Disease, **D)** Ulcerative Colitis, **E)** Schizophrenia, **F)** Body-Mass Index **G)** Waist-Hip Ratio adjusted for body-mass index and **H)** Height. P-values correspond to the association with longitude presented in Table 1.

We applied our approach to five diseases: coronary artery disease (CAD), rheumatoid arthritis (RA), Crohn’s disease (CD), ulcerative colitis (UC) and schizophrenia (SCZ), as well as for three quantitative traits: body-mass index (BMI) waist-hip ratio adjusted for BMI (WHR) and height (HG). We observe that the PS patterns for CAD, RA, SCZ, BMI, HG and WHR closely resemble the main population structure in Finland (Figure 1A). CD and UC do not show clear geographic differences between any parts of the country.

To evaluate statistically whether the PS show geographic differences, we quantified the patterns using a linear model for correlated data, where we explained individuals’ PS with either longitude or latitude and accounted for genetic relatedness of the samples (Methods). The strongest differences were observed for longitude on HG (P = 2.1e-60) and WHR (P = 4.7e-12) and lower but non-zero differences on CAD, RA, SCZ and BMI (all with P<0.05) (Table 1, see Table S1 for results based on the standard linear model without accounting for genetic relatedness). HG and WHR showed differences also for latitude while CD and UC did not show differences either for longitude or latitude. Table 1 also shows that the difference in PS between Eastern Finland (EF) and Western Finland (WF) subpopulations is the largest in HG (-1.51 standard deviation units) and in WHR (1.16). In general, we observed stronger PS differences between east and west than between north and south, which is in line with the main population structure in Finland.

**Table 1.** Results from the linear model for correlated data where polygenic score (PS) is explained by latitude or longitude. SNPs=number of variants in PS. Difference in PS between Eastern Finland (EF) and Western Finland (WF) subpopulations is given in the standard deviation unit of PS. *marks a P-value < 0.05.

| Trait | Study | SNPs | Latitude |  | Longitude |  | EF-WF Difference (95% CI) |
| --- | --- | --- | --- | --- | --- | --- | --- |
|  |  |  | Estimate | P-val | Estimate | P-val |  |
| **CAD** | CARDIoGRAMplusC4D<br>(Nikpey et al. 2015) | 19,597 | -6.3e-4 | 0.97 | 0.05 | 1.6e-4* | 0.63 (0.55, 0.71) |
| **RA** | Okada et al. 2014 | 32,736 | 0.03 | 0.12 | 0.06 | 5.5e-5* | 0.63 (0.55, 0.63) |
| **CD** | IIBDGC<br>(Liu et al. 2015) | 21,771 | 0.03 | 0.18 | -0.002 | 0.87 | -0.10 (-0.19, -0.01) |
| **UC** | IIBDGC<br>(Liu et al. 2015) | 23,513 | 0.03 | 0.23 | 0.02 | 0.22 | 0.26 (0.18, 0.35) |
| **SCZ** | PGC<br>(Ripke et al. 2014) | 30,311 | 0.04 | 8.7e-2 | 0.04 | 4.0e-3* | 0.35 (0.26, 0.43) |
| **BMI** | GIANT<br>(Locke et al. 2015) | 12,742 | 0.03 | 9.4e-2 | 0.04 | 1.8e-3* | 0.53 (0.44, 0.61) |
| **WHR** | GIANT<br>(Shungin et al. 2015) | 13,727 | 0.10 | 1.0e-9* | 0.08 | 4.7e-12* | 1.16 (1.09, 1.23) |
| **HG** | GIANT<br>(Wood et al. 2014) | 27,066 | -0.18 | 1.1e-40* | -0.15 | 2.1e-60* | -1.51 (-1.58, -1.45) |

Recently, it has been reported that PS differences between populations are prone to technical and confounding biases arising especially from population genetic differences (i.e. genetic divergence) or relatedness structure between the GWAS discovery and the target data (Martin et al. 2017; Reisberg et al. 2017; Berg et al. 2018; Curtis 2018; Sohail et al. 2018). To assess whether some of the results in Figure 2 and Table 1 might be affected by these problems, we next concentrate on evaluating our PS in several ways. We use HG as a model trait for developing the methodology.

### Height PS in three independent cohorts

Adult height (HG) is a highly heritable and polygenic trait (Silventoinen et al. 2003; McEvoy and Visscher 2009; Wood et al. 2014), and shows clear phenotypic differences in Finland; Western Finns are on average 1.6 cm taller than Eastern Finns (Methods; Figure 3A). Furthermore, HG is a quantitative trait that makes it possible to compare geographic differences between the observed phenotype and the predictions based on PS. For such comparisons, we regressed out effects of sex, age and age^2^ from HG using residuals from a standard linear model.

**Figure 3.**
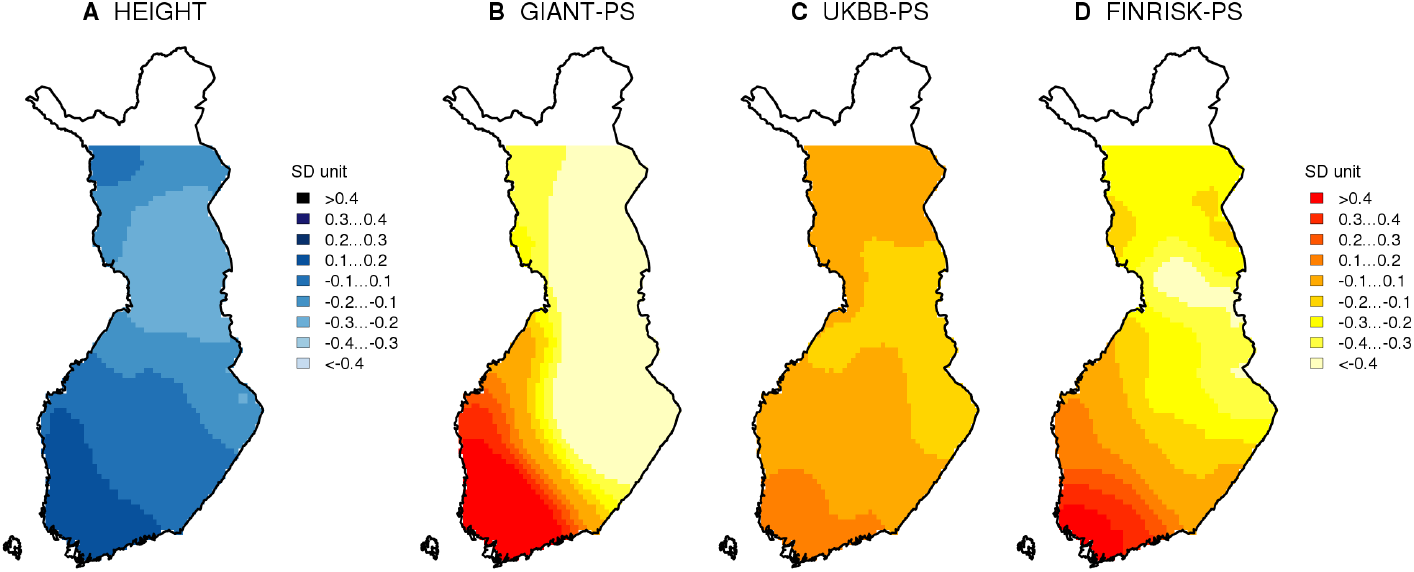
**A)** Distribution of sex, age and age^2^ adjusted adult height and polygenic score (PS) distributions of **B)** GIANT-PS, **C)** UKBB-PS and **D)** FINRISK-PS for height in Finland. The values are in standard deviation unit.

We calculated HG-PS using summary statistics from three independent GWAS, including results from the GIANT consortium (meta-analysis from a heterogeneous set of European samples) (Wood et al. 2014), UK Biobank (single cohort of uniformly genotyped and phenotyped white British samples) and the National FINRISK study (Finnish samples genotyped with two different chips) using our standard pipeline (Methods). Table 2 summarizes the performance of these three scores.

The GIANT consortium height GWAS is a meta-analysis of 250,000 samples from multiple European populations, and it includes about 30,000 Finnish samples (Wood et al. 2014). The GIANT-PS included 27,000 variants, explained 14% of the variance of height and showed dramatic geographic differences in Finland (Figure 3B). The GIANT-PS was 1.5 SD units larger in Western Finland (WF) than in Eastern Finland (EF) and we estimated that this difference would correspond to 3.5 cm predicted height difference between WF and EF by regressing height on this PS in the target sample of 2,376 Finnish individuals (Methods). This difference is over twice the observed phenotypic difference between the subpopulations. Note that even if we assumed that all variation in height were genetic, we would expect our GIANT-PS (that has R^2^ < 15%) to explain only a part of the actual 1.6 cm WF-EF height difference. This raises concerns that GIANT-PS produces geographically biased results in our target sample that cannot be interpreted directly on the phenotypic scale. The predictions were even larger for GIANT-PS if the HG-on-PS regression was done within the WF subpopulation (4.7 cm) or within EF subpopulation (6.4 cm) alone (Table S5), indicating challenges of interpretability for absolute differences among subpopulations.

**Table 2.** Summary of the results in HG-PS comparison. Adjusted R^2^ is the variance explained by the PS in the target set.

| Source |  |  | Finnish | Variants | Adjusted |
| --- | --- | --- | --- | --- | --- |
| GWAS | Ancestry | GWAS N | samples | in PS | $R^2$ |
| GIANT | European | 253,288 | ~35,000 | 27,066 | 14 % |
| UK Biobank | British | 337,199 | 0 | 113,079 | 22 % |
| FINRISK | Finnish | 24,919 | 24,919 | 50,536 | 15 % |

| Source | Observed WF-EF HG-PS | Predicted WF-EF HG |
| --- | --- | --- |
| GWAS | difference (SD unit; 95%CI) | difference (cm; 95%CI) |
| GIANT | 1.51 (1.45, 1.5) | 3.52 (3.14, 3.90) |
| UK Biobank | 0.23 (0.14, 0.32) | 0.64 (0.39, 0.89) |
| FINRISK | 0.59 (0.51, 0.67) | 1.35 (1.14, 1.58) |

Second, we built a PS based on over 330,000 samples of British ancestry from the UK Biobank analyzed by the team led by Benjamin Neale (Churchhouse and Neale 2017). Using the same pipeline as with GIANT-PS, this UKBB-PS contained considerably more variants and gave qualitatively similar geographic results to GIANT-PS but quantitatively showed much smaller WF-EF differences (Figure 3C). UKBB-PS explained 22% of the variation of height in the target sample and corresponded to 0.6 cm predicted WF-EF difference in height.

Third, we built a PS based on the Finnish population-specific summary statistics from the National FINRISK Study (Borodulin et al. 2017). This FINRISK GWAS included nearly 25,000 samples and excluded all our 2,376 target individuals. This FINRISK-PS (50,000 SNPs) explained 15% of the variance of height and showed significant WF-EF differences that corresponded to 1.4 cm difference in predicted height (Figure 3D). For FINRISK-PS and UKBB-PS the predictions were robust to whether the regression was done in the whole target sample or in its WF or EF subset alone (Table S5).

These results show a consistent direction in predicted height differences between Eastern and Western Finland based on three independent GWAS that have different relationships to the target sample. The predicted direction is also consistent with the observed phenotypic difference. However, the results show considerable, and concerning, variation in the predicted geographic difference of genetic component of height.

### Evaluating possible biases in polygenic score for height

#### PS from GIANT accumulates geographic differences

An accumulation of small biases may be a substantial risk in PS of thousands of variants. These small biases can arise, for example, from unadjusted population structure in the underlying GWAS or from overlapping samples between GWAS and target data (Freedman et al. 2004; Marchini et al. 2004). To understand whether the differences in our PS and the unrealistic predictions of geographic height differences may be due to a bias accumulation, we generated additional PS by varying inclusion criteria of variants.

First, we used variants from the initial PS but applied different P-value thresholds. Even though these scores included different numbers of variants, the variance explained did not vary strongly across the thresholds for GIANT-PS or UKBB-PS (Figure 4A), which replicates the behavior of PS reported earlier by Wood et al. 2014 for other target populations. For FINRISK, the variance explained increased considerably with more liberal P-value threshold due to the smaller sample size of the study; specifically, the FINRISK-PS included only a handful of variants for the smallest P-value thresholds and thus had only a little predictive power there (Figure 4A). Conversely, the predicted east-west height differences in the GIANT-PS decreased considerably as more stringent P-value cutoffs were used, while the variance explained for height by the PS increased simultaneously (Figure 4B). The decrease was much subtler for the other two PS. To confirm the effect of the number of variants in the predicted height differences, we randomly sampled 1000 variants from each of the different P-value thresholds in GIANT-PS and calculated the corresponding scores. These PS showed similar levels of predicted WF-EF height difference (about 1 cm) independent of the P-value threshold, which suggests that the number of variants is a more important factor behind the geographic structure of GIANT-PS than the actual phenotypic variance explained by the variants (Figure S1).

**Figure 4.**
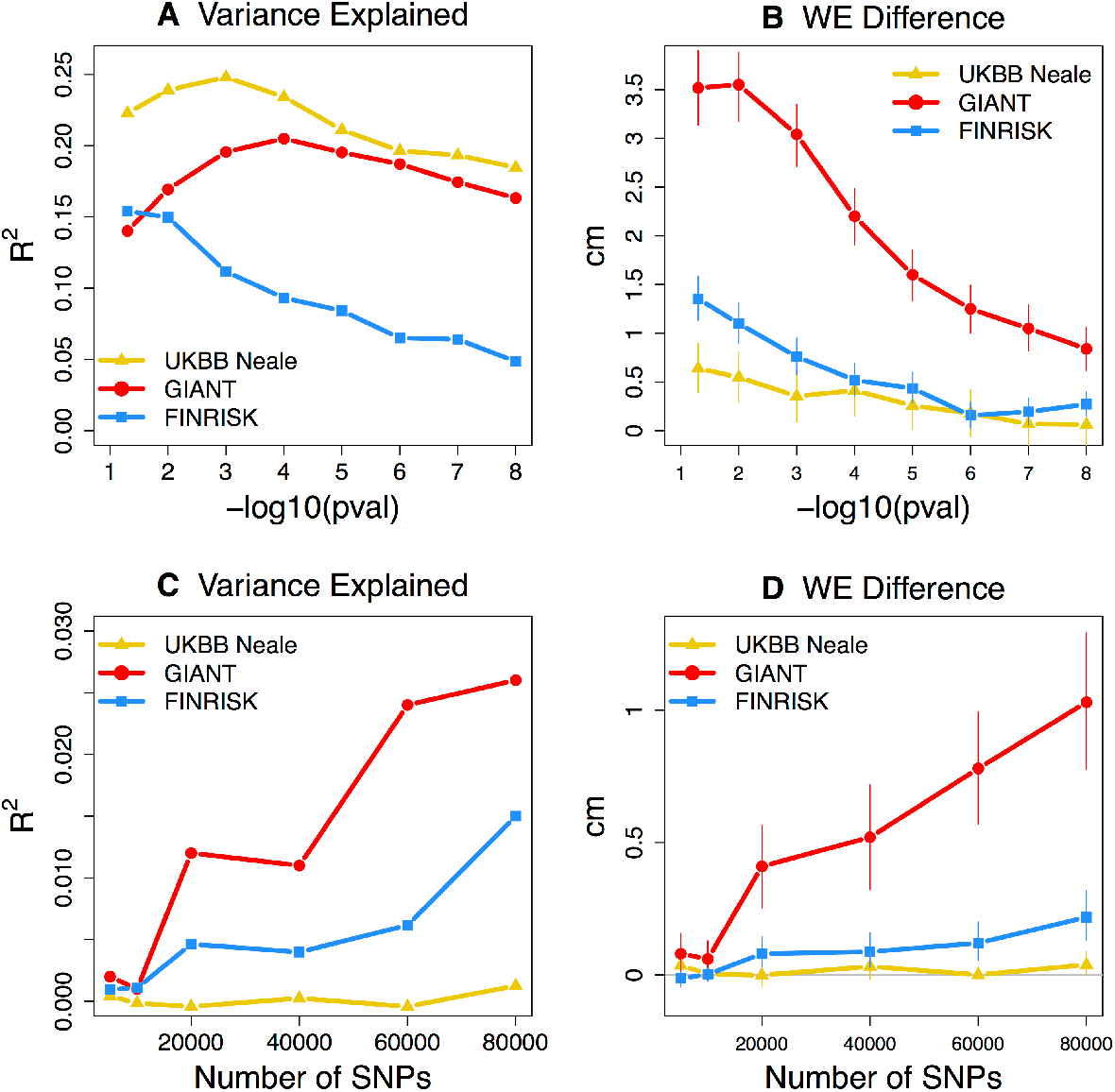
Comparison of PS constructed from height-associated versus random SNPs demonstrates differences in stratification effects by GWAS summary statistics. Top row: **A**) Height variance explained by PS and **B**) predicted East-West difference in height by PS, as a function of P-value threshold in GWAS data. Bottom row: **C**) Height variance explained by PS and **D**) predicted East-West difference in height by PS, as a function of the number of independent variants in PS when all variants have P-value > 0.5 in GWAS. Variance explained is given as adjusted R^2^.

Concerned about the accumulation of WF-EF differences in GIANT-PS, we tested whether similar accumulation occurred even over random, non-associated variants. We randomly sampled different numbers of independent variants whose P-values were over 0.5 in GWAS (suggesting a negligible association to height) and calculated PS for them (we call these “random PS”). This test showed considerable geographic differences in random PS based on GIANT GWAS and these differences increased with the number of variants (Figure 4C-D). Similar but weaker behavior was detected for FINRISK variants but not for UKBB.

One potential explanation for the observed behavior of GIANT-PS is that the effect size estimates have a consistent directional bias that is aligned with the main population structure in Finland. Indeed, GIANT-PS is highly correlated with the leading principal component (PC1) of population structure in our target sample (r = 0.80) and when we removed the linear effect of PC1 from the GIANT-PS (Methods), the residuals explained more variance (19%) of height than the original GIANT-PS (14%). This suggests that in the effect size estimates of GIANT-PS, a part of the true height association is masked by a strong component aligned with PC1 in Finland. For neither FINRISK-PS nor UKBB-PS did the removal of the linear effect of PC1 improve the variance explained in our target sample (Table S2). To further test the possible bias in GIANT effect estimates, we took the overlapping HG variants between the GIANT-PS and the UKBB data (overlap was 26,853 out of 27,066 variants) and made a new GIANT-UKBB-PS (i.e. using GIANT variants but UKBB effects). We observed that this GIANT-UKBB-PS explained 26% of the height variance in the target sample and, contrary to GIANT-PS, its predictive power was not masked by PC1 (R^2^ dropped from 26% to 24% after regressing out PC1 from GIANT-UKBB-PS). Multiple regression unequivocally showed that GIANT-PS was not predictive of HG (P = 0.25) when GIANT-UKBB-PS was simultaneously included in the model (P = 3e-84). These observations confirm that while the variants in GIANT-PS are useful for predicting HG in Finnish samples, their estimates reported in GIANT are unrealistically strongly associated with the population structure in Finland and that the UKBB effect size estimates considerably improve the predictive power of the corresponding PS. However, we also observed that, despite more accurate effect estimates, the GIANT-UKBB-PS predicted surprisingly large WF-EF difference (2.5 cm [2.0, 2.8]). This suggests that it is not only the bias in effect estimates that drives the geographic difference but also the choice of the variants. We speculate that the variant that has the smallest HG P-value in GIANT in any one genomic region tends to be one of the most geographically stratified variants among all HG associated variants in that region. This could lead to a PS that is enriched with geographically stratified variants and therefore emphasize geographic differences in PS even when the effect estimates were unbiased for HG.

Together, these analyses suggest that the geographic distribution of PS based on the GIANT summary statistics consistently exaggerates height differences between the main Finnish subpopulations, whereas much less confounding from population stratification is seen in FINRISK-PS, and almost none is observed in UKBB-PS. A few possible reasons for this bias accumulation could be inadequate adjustment for population structure in GWAS (Sohail et al. 2018, Berg et al. 2018) or partially overlapping or related samples between GWAS samples and test data. Next, we consider the effect of overlapping and related samples.

#### Effect of overlapping samples

Our target data originate from the National FINRISK Study that is not reported among the GIANT cohorts (neither among 46 cohorts in (Lango Allen et al. 2010) nor among additional 32 cohorts in Wood et al. 2014). However, a closer look into the cohort descriptions suggested that COROGENE, DILGOM, MORGAM and MIGEN cohorts may include some FINRISK samples. This shows that it is not always straightforward to keep track of where publicly accessible samples have been used previously, which would be a crucial piece of information for appropriately validating PS. While computational methods exist to detect sample overlaps between GWAS summary statistics (Bulik-Sullivan et al. 2015), their behavior in large datasets are not yet completely understood (Yengo et al. 2018).

To test whether overlapping individuals affect results, we ran an additional GWAS on HG using the FINRISK samples but now also included the 2,376 target individuals in the GWAS. We then built a PS (called FINRISK-OV-PS for FINRISK-OVERLAP-PS) based on this GWAS and compared it to our original FINRISK-PS where the target individuals were excluded. FINRISK-OV-PS was naturally overfitted to the target data and consequently explained 74% of the height variance in the target sample. Despite this overfitting, it predicted only a 1.5 cm [95% CI: 1.07 – 1.93] height difference between WF and EF, which is very similar to the prediction of our original FINRISK-PS (1.4 cm [95% CI: 1.14 – 1.58]). This demonstrates that even a large number of overlapping samples may have only a small effect on the predicted geographic difference between the populations even though the overfitting effect on the variance explained may be enormous. Thus, the possible overlap between GIANT GWAS and some of our target individuals is unlikely to be the main reason for the large difference in geographic distribution between GIANT-PS and the other PS.

#### Effect of related individuals and residual population structure

Cryptic relatedness or unadjusted population structure can cause bias in GWAS that can affect the PS. We assessed the effects of these factors by comparing our initial HG-PS based on the results from the GWAS of standard linear regression using principal component adjustment to a PS based on the GWAS using a linear mixed model implemented in BOLT-LMM (Loh et al. 2015), both in FINRISK and UKBB data. We could not apply the mixed model to GIANT data since the individual-level data were not available to us. Results (Table S3) show that for UKBB, the PS based on the mixed model explained slightly more variance of height (25%) in our target sample than the PS based on linear regression (22%), while for FINRISK, the variance explained remained the same (15%). The predicted height differences between WF and EF based on the BOLT-LMM summary statistics were 0.37 cm (0.10 – 0.64 cm) for UKBB and 1.15 cm (0.94 – 1.38 cm) for FINRISK. Compared to the standard linear regression, 0.64 cm (0.39 – 0.89 cm) for UKBB and 1.35 cm (1.14 – 1.58 cm) for FINRISK, PS based on the mixed model gave smaller predicted differences, although the differences between the mixed model and the standard linear model were not statistically significant. This suggests that there can be some residual population structure and/or cryptic relatedness using standard linear regression with the leading principal components as covariates, resulting in larger geographical differences in the PS. We further tested whether the mixed model reduced the accumulation of population differences in the random PS in FINRISK data but the accumulation remained similar to the linear model results (Figure S2). Another way how cryptic relatedness could affect PS is through non-random relatedness between the GWAS and target samples. Later in this work, we will use SCZ data to test whether including some Finnish samples in GWAS has an effect on geographic differences of SCZ PS.

### Testing bias accumulation in other complex diseases and traits

#### Accumulation of geographic differences in other diseases and traits

After assessing multiple sources of bias accumulation in HG, we applied similar strategies to the other seven phenotypes. For each phenotype, we generated random PS with increasing number of variants to detect a possible accumulation of biases. Here we present absolute difference between Eastern and Western Finland using standardized PS since we did not have a way to turn these to the phenotypic scale for disease studies. Figure 5 shows that for RA, CD, UC and SCZ, the absolute WF-EF PS difference of the random score is close to zero whereas for CAD, BMI, WHR and HG we observe a possible accumulation of bias.

**Figure 5.**
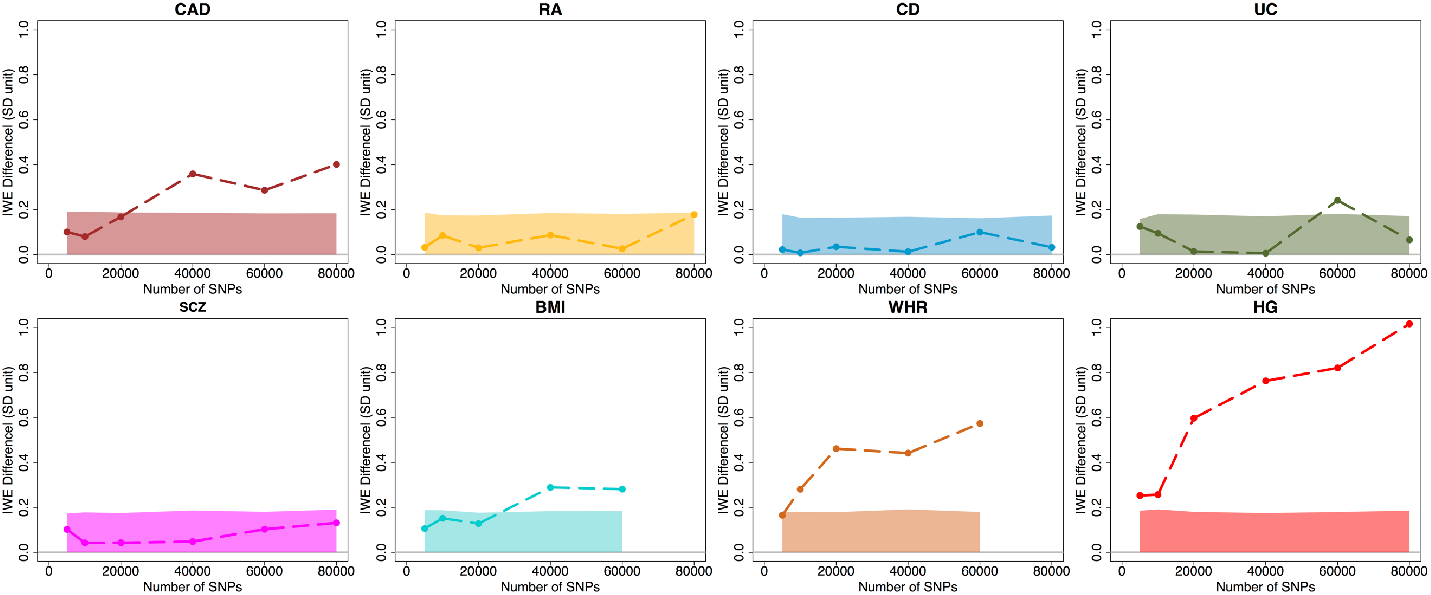
Absolute value of polygenic score difference between Eastern and Western subpopulations using different numbers of independent variants (r^2^ < 0.1) randomly chosen with GWAS P-value > 0.5. For BMI and WHR, the data did not contain more than 60,000 independent variants. The solid region is the 95% probability interval under the theoretical null assumption of zero effect sizes and completely independent variants (r^2^ = 0) (Methods).

Similarly to HG, we were able to compare the geographic distribution of PS of BMI and WHR to their phenotypic counterparts. Figures S3A and S6A show that neither BMI nor WHR shows clear geographic patterns in our data. For BMI, the comparison of PS based on three different GWAS (Figure S3 B-D and Figure S4) shows that GIANT-PS has the largest variance explained (8.0%) and the largest geographic difference and also shows signs of an accumulation of geographic bias between populations when we use PS based on a random set of non-associated variants. UKBB-PS does not show any evidence of EF-WF difference matching the observed phenotypic distribution (Figure S3) and explains 4.9% of variance. FINRISK-PS explains the least amount of phenotypic variance (1.3%) and shows only a limited amount of EF-WF difference. For WHR, GIANT-PS explained 2.0% of variance and showed again a dramatic geographic difference both for the initial PS and a random set of non-associated variants. FINRISK-PS explained 1.1% of variance and showed a very subtle difference between East and West (Figures S5, S6 and Table S7). The UKBB GWAS results available at Neale lab summary statistic resource did not contain WHR. These results on BMI and WHR largely repeat the results on HG with respect to the behavior of PS as a function of the three different GWAS cohorts.

#### Effect of Finnish samples in schizophrenia

Table 3 summarizes the properties of GWAS that we used. It is noteworthy that all phenotypes with Finnish samples included in the GWAS also showed some bias accumulation in our analyses (Figure 5). Because Finns are considered a genetic isolate (Lappalainen et al. 2006; Jakkula et al. 2008; Kerminen et al. 2017; Martin et al. 2018b), we tested whether including a small number of Finnish samples (but not our target individuals) had an effect on geographic distribution of PS. SCZ was the only phenotype in Table 3 where we had access to the original GWAS data. Summary statistics of SCZ reported by Psychiatric Genomics Consortium (Schizophrenia Working Group of the Psychiatric Genomics 2014) included 546 Finnish cases and 2,011 Finnish controls, while in our analyses so far, we have used summary statistics where these individuals were excluded. We compared the results of the PS based on the GWAS excluding (SCZ-PS) and including the Finnish samples (SCZ-INCL-PS) and, overall, the genetic risk maps show similar risk patterns (Figure S7). Quantitatively, the geographic difference in the PS difference is stronger in SCZ-INCL-PS (Table S4), which varies significantly both in the east-west (P < 3e-4) and north-south (P < 4e-6) directions, while SCZ-PS shows significant differences only in the east-west (P < 4e-3) direction.

**Table 3.**
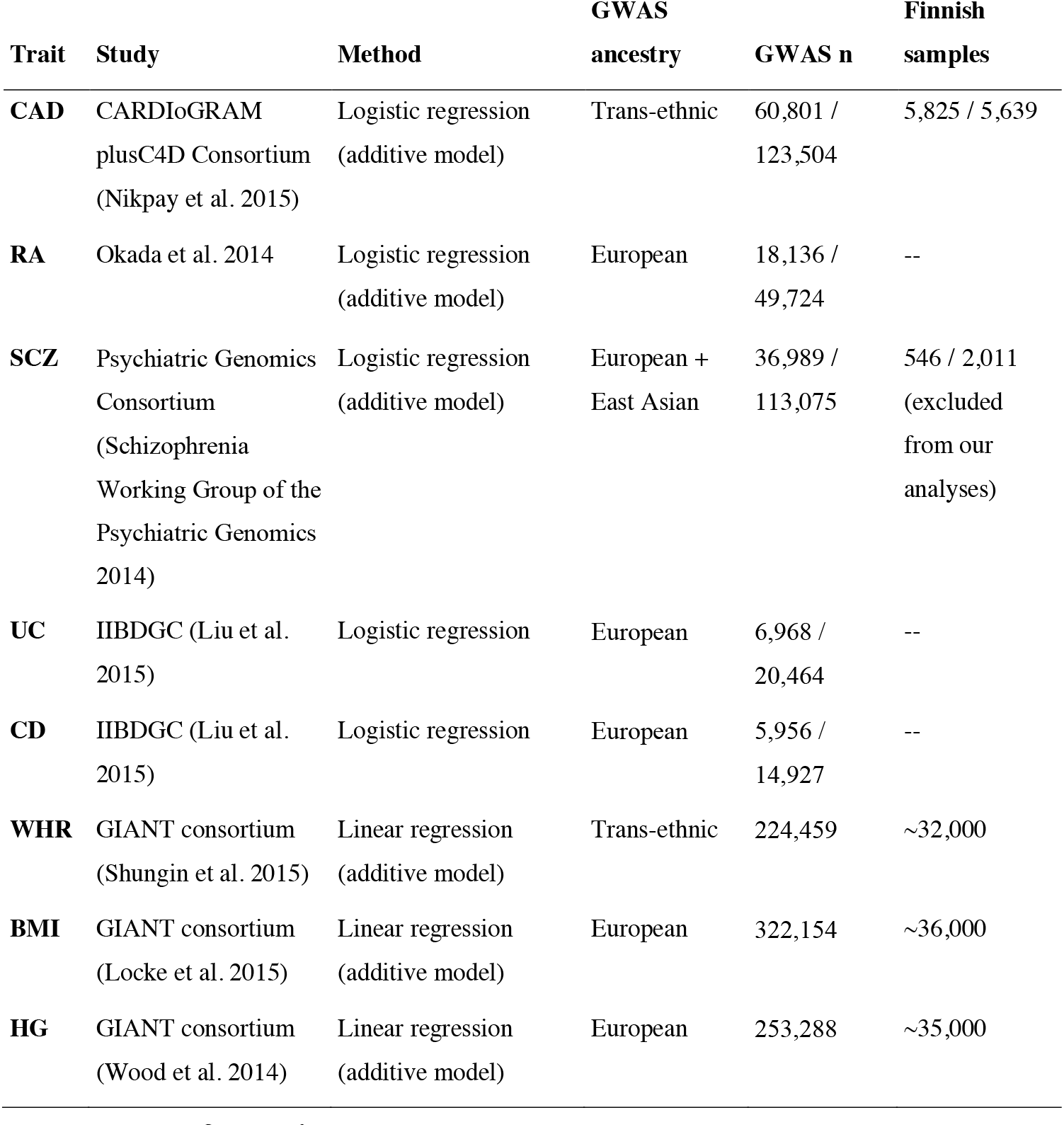
Summary of the GWAS statistics and sources of possible bias. For diseases, GWAS n = cases / controls.

Thus, there is a possible effect of cryptic relatedness between the original GWAS sample and our target sample. However, since this effect seems to be stronger in north-south direction, we do not observe an accumulation of differences over randomly sampled variants between the EF and WF populations (Figure S8).

## Discussion

Polygenic scores (PS) have recently reached predictive power of some well-established monogenic risk factors for disease (Khera et al. 2018), and several projects are currently testing their utility in health care settings. PS could also potentially inform us about the role of genetics in geographic variability of traits and disease. However, a major challenge is that the geographic distribution of PS is a complex function of population genetic differences between the GWAS data and the target samples, complicating its interpretation (Scutari et al. 2016; Martin et al. 2017; Reisberg et al. 2017; Martin et al. 2018a). Here we studied the geographic distribution of several PS within Finland and assessed their robustness and possible biases in several ways.

By generating PS for eight phenotypes on Finnish samples, we observed strong similarities between the geographic distribution of several PS and the main population structure in Finland that runs from south-west to north-east (Kerminen et al. 2017). We further showed that even the least statistically significantly associated half of the effect sizes (with GWAS P-value > 0.5) were carrying a consistent pattern of east-west difference for CAD (CARDIoGRAM data) and the three anthropometric traits from the GIANT consortium: HG, BMI and WHR, which we interpret indicating a likely bias. In theory, such a pattern could also result from extreme polygenicity. However, with the highly polygenic HG as our model trait, we showed that the random score from our largest HG GWAS based on the UK Biobank did not show any east-west variation within Finland. This suggests that the geographic difference accumulating in the random score from GIANT is rather due to bias than polygenicity. Furthermore, we observed for HG that the GIANT-PS was so strongly aligned with the first principal component of the genetic structure in our target data that this association masked some of the predictive power of the PS. This suggests that the effect estimates from GIANT contains a bias aligned with the main population structure in Finland, which is in line with two recent studies that have reported related biases in the context of polygenic selection studies (Berg et al. 2018; Sohail et al. 2018).

For all three quantitative traits, PS predicted unrealistically large geographic differences compared to the actual phenotypic differences. A theoretical but unlikely possibility remains that the geographic structure of the genetic component not explained by our current PS could be opposite to the component that is explained by our current PS, which could eventually balance out the unrealistically large estimates for GIANT-PS and FINRISK-PS. However, given that the estimated difference consistently increases with the inclusion of more variants in PS, a more plausible explanation is that we simply cannot robustly interpret the geographic differences in PS derived from existing GWAS on the phenotypic scale via a simple regression framework. Earlier, results of phenotypically inconsistent PS differences between continental groups have been reported (Martin et al. 2017; Reisberg et al. 2017; Curtis 2018). Here we show that similar patterns can exist even for a relatively small geographic area and a relatively homogeneous population of Finland. We note that even if the genetic E-W difference in Finland may be large compared to variation within some other European countries (Salmela et al. 2008), it is tiny compared to the continental differences (The 1000 Genomes Project Consortium 2012).

Our results showed that the phenotypes that did not accumulate E-W differences were the two types of inflammatory bowel disease (CD and UC), and SCZ and RA. Of these, CD and UC did not show any geographic PS variation in Finland. To our knowledge, only two studies have studied the geographic variation in the prevalence of inflammatory bowel disease (IBD) in Finland. Lehtinen et al. 2016 reported higher incidence rates of pediatric IBD in more sparsely populated areas while Jussila et al. 2013 reported increasing UC prevalence rates in Northern Finland but no geographic structure for CD. Our polygenic risk prediction for CD is in line with the observations in Jussila et al. (2013) and, even though the PS of UC did not show significant geographic differences in our statistical analysis, the genetic risk map for UC shows some increasing risk pattern in Northern Finland. SCZ showed a higher polygenic risk in EF than in WF, which is in line with extensive geographic incidence information from several studies (Lehtinen et al. 1990; Hovatta et al. 1997; Haukka et al. 2001; Perala et al. 2008; Pietiläinen 2014; Kurki et al. 2018) that describe highest SCZ prevalence/incidence rates in Northern and Eastern Finland and lowest rates in the Southwestern parts of the country. Also, RA showed higher polygenic risk in EF than in WF. Our limited information about regional incidence of RA in Finland is from (Kaipiainen-Seppanen et al. 2001) who reported highest RA incidence rates for North Karelia (in EF) and lowest for Ostrobothnia (on the west coast), but unfortunately the study did not include Southwestern or Northern Finland.

Neither our SCZ nor our RA GWAS summary statistics included any Finnish samples. Together these two diseases exemplify the potential of PS to explain geographic health differences.

To conclude, we recommend the following practices for geographic evaluation of PS. (1) Check residual geographic stratification of PS by generating random scores of non-associated variants and by testing whether PS unrealistically strongly align with the leading PCs of genetic structure. (2) Use a linear/logistic mixed model instead of the standard linear/logistic regression model in GWAS. (3) Compare the genetically predicted phenotypic difference between populations to the observed phenotypic difference to detect unrealistic genetic predictions. With these tools, we showed that while PS for several traits in Finland followed the geographic distribution of the phenotype (HG, CAD, SCZ, RA, CD, UC), for CAD, HG as well as for BMI and WHR, we observed suspicious behavior of the geographic distribution of PS that could indicate a bias arising from population genetic structure rather than from a direct genotype-phenotype association. Our results emphasize that we have limited understanding of the interplay between our current PS and genetic population structure even within one of the most thoroughly studied populations in human genetics. Therefore, we recommend refraining from using the current PS to argue for significant polygenic basis for geographic phenotype differences until we understand better the source and extent of the geographic bias in the current PS.

## Materials and Methods

### Geographically defined target data

We used data from the National FINRISK Study which is a survey for Finnish adult population (age from 25 to 74) to estimate risk and protecting factors of chronic diseases (Borodulin et al. 2017). The FINRISK Study has collected several thousand samples every five years, since 1972. We used data from the FINRISK Study survey of 1997 on geographically defined sample of 2,376 individuals that was previously described in Kerminen et al. 2017. The two parents of each individual in this sample were both born within 80 km from each other. For the genetic analyses, we used genotypes from Illumina HumanCoreExome-12 BeadChip (see details in Kerminen et al. 2017) and imputed genotypes as described in (Ripatti et al. 2016).

### Variant filtering for polygenic scores

We derived polygenic scores (PS) in our target data for each disease and trait based on large international GWAS meta-analyses whose summary statistics were publicly available. We derived all PS by excluding variants whose minor allele frequency (MAF) was below 1% in meta-analysis or whose meta-analysis P-value was above 0.05 or that resided in the major histology complex (chr 6: 25-34 Mb) (Price et al. 2008). In addition, where applicable, we filtered out variants whose INFO score was below 90% or that had been present in less than 90% of the cohorts of the meta-analysis. We also excluded all multi-allelic variants. Finally, the PS were built by selecting independent variants with PLINK 1.9 (Purcell and Chang; Chang et al. 2015), using clump command with 500 kb window radius and 0.1 threshold for r^2^. Number of remaining variants in each PS are given in Tables 1 and 2. Below, we give the detailed information of the data and filtering for each disease and trait separately.

#### CAD

The CAD-PS was derived from the CARDIoGRAMplusC4D study (Nikpay et al. 2015) and the summary statistics were based on the cohort with trans-ethnic ancestry. Variant filtering was based on above MAF and P-value thresholds. INFO score filtering was used. Variant was also filtered out if it was not observed in at least 90% of the cohorts in meta-analysis.

#### RA

The RA-PS was derived from study of Okada et al. (2013) and the summary statistics were based on the cohort with European ancestry. Variant filtering was based on above MAF and P-value thresholds. INFO score filtering was not used because it was not available. MAF was based on the FINRISK data. Filtering based on the number of cohorts or samples was not used.

#### CD

The CD-PS was derived from the International Inflammatory Bowel Disease Genetics Consortium study (Liu et al. 2015) and the summary statistics were based on the cohort with European ancestry. Variant filtering was based on above MAF and P-value thresholds. INFO score filtering was used. Variant was also filtered out if it was not observed in at least 90% of the cohorts in meta-analysis.

#### UC

The UC-PS was derived from International Inflammatory Bowel Disease Genetics Consortium study (Liu et al. 2015) and the summary statistics were based on the cohort with European ancestry. Variant filtering was based on above MAF and P-value thresholds. INFO score filtering was used. Variant was also filtered out if it was not observed in at least 90% of the cohorts in metaanalysis.

#### SCZ

The SCZ-PS was derived from Psychiatric Genomics Consortium study (Schizophrenia Working Group of the Psychiatric Genomics 2014) and the summary statistics were based on the cohort with European and Asian ancestry from which the Finnish samples (546 cases and 2,011 controls) were excluded. Variant filtering was based on above MAF and P-value thresholds. INFO score filtering was used. Variant was also filtered out if it was not observed in at least 90% of the cohorts in metaanalysis.

#### HG

The HG-GIANT-PS was derived from the GIANT consortium study (Wood et al. 2014) and the summary statistics were based on the sex-combined cohort with European ancestry. Variant filtering was based on above MAF and P-value thresholds. INFO score filtering was not used as it was not available. Variant was also filtered out if it was not observed in at least 90% of samples. The HG-UKBB-PS was derived from the UK Biobank data analyzed by the Neale lab (Churchhouse and Neale 2017) and the summary statistics were based on the cohort with white-British ancestry. Variant filtering was based on above MAF and P-value thresholds. INFO score filtering was not used as it was not available. Filtering based on the number of cohorts or samples was not used.

The HG-FINRISK-PS was derived from the National FINRISK Study (Borodulin et al. 2017) and the summary statistics were based on the cohort with Finnish ancestry (see details below). Variant filtering was based on above MAF and P-value thresholds. INFO score filtering was not used. Filtering based on the number of cohorts or samples was not used.

#### BMI

The BMI-GIANT-PS was derived from the GIANT consortium study (Locke et al. 2015) and the summary statistics were based on the sex-combined cohort with European ancestry. Variant filtering was based on above MAF and P-value thresholds. INFO score filtering was not used as it was not available. Variant was also filtered out if it was not observed in at least 90% of samples.

The BMI-UKBB-PS was derived from the UK Biobank data analyzed by the Neale lab (Churchhouse and Neale 2017) and the summary statistics were based on the cohort with white-British ancestry. Variant filtering was based on above MAF and P-value thresholds. INFO score filtering was not used as it was not available. Filtering based on the number of cohorts or samples was not used.

The BMI-FINRISK-PS was derived from the National FINRISK Study (Borodulin et al. 2017) and the summary statistics were based on the cohort with Finnish ancestry (see details below). Variant filtering was based on above MAF and P-value thresholds. INFO score filtering was not used. Filtering based on the number of cohorts or samples was not used.

#### WHR

We considered BMI-adjusted WHR throughout this work.

The WHR-GIANT-PS was derived from the GIANT consortium study (Shungin et al. 2015) and the summary statistics were based on the sex-combined cohort with European ancestry. Variant filtering was based on above MAF and P-value thresholds. INFO score filtering was not used as it was not available. Variant was also filtered out if it was not observed in at least 90% of samples.

The WHR-FINRISK-PS was derived from the National FINRISK Study (Borodulin et al. 2017) and the summary statistics were based on cohort with Finnish ancestry (see details below). Variant filtering was based on above MAF and P-value thresholds. INFO score filtering was not used. Filtering based on the number of cohorts or samples was not used.

### Additional GWAS for FINRISK and UK Biobank

#### FINRISK

We ran two standard linear regressions for HG, BMI and WHR (adjusted for BMI) first using 27,294 individuals across the National FINRISK Study collections 1992-2012 and second excluding all our target individuals from the first set. The linear regression was done in HAIL (Hail). Both GWAS used sex, age, FINRISK project year, genotyping chip and the first 10 principal components of population structure as covariates in the analysis. In addition, we ran a linear mixed model for the second data set with the same covariates as with the standard linear model using BOLT-LMM v.2.3 (Loh et al. 2015).

#### UKBB

For the UK biobank, we performed a linear mixed model GWAS for HG with BOLT-LMM v2.3 (Loh et al. 2015). For this analysis, we mimicked the linear regression analysis (round 1) of the Neale lab (Churchhouse and Neale 2017) and used UKBB v2 genotypes on 343,728 samples with white-British ancestry. We used age, sex and the first 20 principal components as covariates and used directly genotyped variants with MAF above 1% and missingness below 10% for generating the variance component. GWAS statistics were calculated for imputed data with MAF above 0.1% and INFO above 0.7.

### Polygenic scores

We calculated polygenic scores for the target set of 2,376 FINRISK individuals using the additive model as

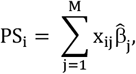

where *PS_i_* is a polygenic score for individual *i*, M is the number of SNPs in the score (after variant filtering), *x_ij_* is the individual’s (imputed) genotype dosage for SNP *j*, and 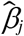 is the effect size estimate of SNP *j* from the GWAS.

### Genetic risk maps

To visualize the geographic distribution of PS, we used geographic locations of our geographically well-defined sample of 2,376 individuals and their PS. We estimated individual’s geographic location as the mean of his/her parents’ birth places. Risk maps were then created in R using a geographical centroid approach: it lays a grid on the map of Finland and for each grid point *p* calculates the average of individuals’ PS inversely weighted by their squared distance to the grid point as

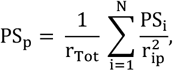

where *r_ip_* is the distance between individual *i* and grid point *p*, and 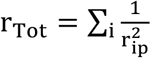 is the sum of the weights. We used a grid with a square size of 10 kilometers and limited the minimum value for *r_ip_* to be 50 kilometers to avoid high variance in weights. In addition, to control for uncertainty in the areas that have a low sample size, we added to the calculation of *PS_p_* one pseudo-individual whose PS is the population average PS and whose distance to the point *p* is the minimum of the observed distances *r_ip_*. This modification draws the PS values of grid points towards the population average especially at sparse areas where there are few individuals at the minimum range from the grid point. Last, the risk maps were scaled by the population average and standard deviation using a subset of 1,042 geographically evenly distributed individuals as described in Kerminen et al. (2017). The border line for the map of Finland was obtained from https://gadm.org/.

### Linear model for correlated data to assess spatial PS differences

To quantify whether the polygenic score has geographic differences, we performed a regression analysis using a linear model for correlated data where we explained PS with latitudinal or longitudinal coordinate and accounted for genetic relatedness as

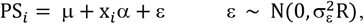

where *x_i_* is the coordinate of individual *i*, μ is the intercept and a is the effect of latitude or longitude on PS reported in Tables 1, S4, S6 and S7. For the structure of the error terms, we used the genetic relationship matrix *R* that was estimated with PLINK 1.9 (Purcell and Chang; Yang et al. 2011; Chang et al. 2015) (command --make-rel) using 61,598 independent variants from Illumina HumanCoreExome chip described in Kerminen et al. (2017). Regression results with the standard linear model without accounting for genetic relatedness are shown in Table S1.

### Polygenic and phenotypic differences between subpopulations

The two main subpopulations in Finland are located in Eastern Finland (EF) and Western Finland (WF) and were previously described in Kerminen et al. (2017) and shown in Figure 1A. Here, we reproduced this analysis using CHROMOPAINTER and FineSTRUCTURE (Lawson et al. 2012) with our current sample of 2,376 individuals to estimate both phenotypic and polygenic score differences between these two populations. The analysis divided our target sample into 1,604 EF and 772 WF individuals and this division was used for estimating the differences between subpopulations.

#### PS differences in standard deviation units

We calculated the PS differences between the subpopulations by first scaling the PS of the target sample with the subset of geographically evenly distributed 1,042 samples. Scaled PS were then used to calculate the difference between EF and WF. This strategy ensured a robust comparison between PS based on a fixed reference set. The 95% confidence intervals for the difference between two groups were given by Welch’s t-test in R 3.4.1 (R Core Team 2018).

#### Phenotypic differences predicted by PS

For HG, BMI and WHR, we estimated also the phenotypic difference predicted by PS between our subpopulations. First, we fitted the linear model where we explained the phenotype with general covariates sex, age and age^2^ (WHR was additionally adjusted for BMI) in our target sample and then we fitted another linear model where we explained the residuals with PS. Based on the effect estimates of the second model, we were able to estimate the predicted phenotypic effect by multiplying the PS effect estimate with the PS difference between the populations. We estimated the respective 95% credible intervals by simulation approach where we generated 100,000 samples of pairs of effect estimates for PS difference d and PS effect on phenotype β from their posterior distributions (assuming improper flat priors) and by using the empirical distribution of d*β as the posterior distribution. The posterior distribution of d was modeled as a Normal distribution with mean set to the observed PS difference and standard deviation calculated from the 95% confidence interval from the Welch’s t-test as 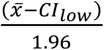. The posterior for β was modeled as a Normal distribution with mean set to the observed effect estimate and standard deviation set to the corresponding standard error from the linear model.

#### Observed phenotypic differences for HG, BMI and WHR

We estimated the observed phenotypic difference in HG, BMI and WHR between EF and WF by adjusting the corresponding trait for sex, age and age^2^ (WHR was additionally adjusted for BMI) using linear regression and then calculated the difference of the subpopulation means based on the residuals from this regression. The residuals were maintained in the units of the original phenotypes.

### P-value thresholding in PS

We studied the effect of P-value threshold for our PS by applying 7 different thresholds (P-value < 1e-2, 1e-3, 1e-4, 1e-5, 1e-6, 1e-7 and 1e-8) to the variants of initial PS (that used threshold 0.05) and calculated the additional PS as described above.

### Accumulation of biases using weakly associated variants (“random PS”)

To detect accumulation of biases we used a simple approach where we first filtered the GWAS summary statistics similarly to the original scores (as explained above) except that we considered only variants with the GWAS P-value larger than 0.5. This left us with at most very weakly associated variants. Among these variants, we preformed LD clumping using same parameters as above except that we set the P-value cut-off to 1 in order to not further exclude any variant based on its P-value and we permuted the P-values among the variants to ensure that the resulting scores are random with respect to their P-value. LD clumping resulted in different number of variants for different traits and from those we randomly sampled increasing number of variants (5,000; 10,000; 20,000; 40,000; 60,000; 80,000). For BMI and WHR the remaining number of variants after LD clumping was < 80,000 and hence we were not able to compute a PS for 80,000 for these two traits. Finally, we calculated the PS for each individual and evaluated the difference between subpopulations in these “random PS”.

To understand the expected behavior of PS with truly zero effect sizes and to compare with our observed random PS, we generated 1,000 simulated PS for each observed random PS. These PS were simulated using the variants from the random PS but sampling their effect estimates independently from a normal distribution with mean zero and standard deviation that corresponded to the standard error of the variant in GWAS. In Figure 5 we see that the 95% highest probability interval of the population difference is approximately constant across the different number of variants in the PS and across the different GWAS. Supplemental text describes the theoretical basis for this property.

Our simulated 95% intervals assume completely independent variants whereas our PS pipeline used a more liberal LD threshold of r^2^ < 0.1. Therefore, we also compared the effects of residual LD to our random scores by performing LD-clumping with r^2^ threshold of 0.01 and 0.001 for CAD and HG with GIANT and UKBB data. The Supplemental Figure S9 shows that, for GIANT-PS, the residual LD does not have an effect on the accumulation of population difference and similar tendency is suggested for CAD-PS even though the data are limited. For UKBB-PS, there is no accumulation of difference for any r^2^ threshold.

## Supporting information

## Acknowledgments

We thank the participants of the FINRISK cohort and its funders: the National Institute for Health and Welfare, the Academy of Finland (139635 to VS), and the Finnish Foundation for Cardiovascular Research. This research was conducted with the UK Biobank Resource under application no. 22627. Data on coronary artery disease have been contributed by CARDIoGRAMplusC4D investigators and have been downloaded from www.CARDIOGRAMPLUSC4D.ORG. We thank PGC Schizophrenia group, IBDGC, GIANT and RA consortia and the Neale lab for providing summary statistics. This work was financially supported by the University of Helsinki Doctoral Programme in Population Health (SK), the Academy of Finland (288509, and 294050 to MPi; 251217 and 285380 to SR) and its Center of Excellence in Complex Disease Genetics (1312076 to MP; 213506 and 129680 to SR) and by the Research Funds of the University of Helsinki to MPi. SR was further supported by EU FP7 projects ENGAGE (201413), BioSHaRE (261433), the Finnish Foundation for Cardiovascular Research, Biocentrum Helsinki, and the Sigrid Jusélius Foundation. ARM was supported by funding from the National Institutes of Health (K99MH117229).

## Author contributions

SK, SR and MPi designed the study. SK conducted the analyses. JK and SER conducted the genome-wide association studies on FINRISK data. MPi derived the mathematical formulas. ASH, MPe and VS provided materials. IS provided methods. SK, ARM, AP, MJD, SR and MPi interpreted the results. SK and MPi wrote the manuscript with help from ARM. All authors reviewed the manuscript.

## References

Abraham, G., A.S. Havulinna, O.G. Bhalala, S.G. Byars, A.M. De Livera et al., 2016 Genomic prediction of coronary heart disease. Eur Heart J 37 (43):3267–3278.

Berg, J.J., A. Harpak, N. Sinnott-Armstrong, A.M. Joergensen, H. Mostafavi et al., 2018 Reduced signal for polygenic adaptation of height in UK Biobank. bioRxiv, https://doi.org/10.1101/354951

Borodulin, K., H. Tolonen, P. Jousilahti, A. Jula, A. Juolevi et al., 2017 Cohort Profile: The National FINRISK Study. Int J Epidemiol. 47 (3):696–696i.

Bulik-Sullivan, B.K., P.R. Loh, H.K. Finucane, S. Ripke, J. Yang et al., 2015 LD Score regression distinguishes confounding from polygenicity in genome-wide association studies. Nat Genet 47 (3):291–295.

Chang, C.C., C.C. Chow, L.C. Tellier, S. Vattikuti, S.M. Purcell et al., 2015 Second-generation PLINK: rising to the challenge of larger and richer datasets. Gigascience 4:7.

Churchhouse, C., and B.M. Neale, 2017 Rapid GWAS of thousands of phenotypes for 337,000 samples in the UK Biobank. http://www.nealelab.is/blog/2017/7/19/rapid-gwas-of-thousands-of-phenotypes-for-337000-samples-in-the-uk-biobank

GBD 2016 Healthcare Access Quality Collaborators, 2018 Measuring performance on the Healthcare Access and Quality Index for 195 countries and territories and selected subnational locations: a systematic analysis from the Global Burden of Disease Study 2016. Lancet 391 (10136):2236–2271.

The 1000 Genomes Project Consortium, 2012 An integrated map of genetic variation from 1,092 human genomes. Nature 491 (7422):56–65.

Curtis, D., 2018 Polygenic risk score for schizophrenia is more strongly associated with ancestry than with schizophrenia. Psychiatr Genet 28 (5):85–89.

The Finnish Disease Heritage. http://www.findis.org/heritage.html

Freedman, M.L., D. Reich, K.L. Penney, G.J. McDonald, A.A. Mignault et al., 2004 Assessing the impact of population stratification on genetic association studies. Nat Genet 36 (4):388–393.

Fuchsberger, C., J. Flannick, T.M. Teslovich, A. Mahajan, V. Agarwala et al., 2016 The genetic architecture of type 2 diabetes. Nature 536 (7614):41–47.

Hail. https://github.com/hail-is/hail

Hartiala, J., W.S. Schwartzman, J. Gabbay, A. Ghazalpour, B.J. Bennett et al., 2017 The Genetic Architecture of Coronary Artery Disease: Current Knowledge and Future Opportunities. Curr Atheroscler Rep 19 (2): 6.

Haukka, J., J. Suvisaari, T. Varilo, and J. Lonnqvist, 2001 Regional variation in the incidence of schizophrenia in Finland: a study of birth cohorts born from 1950 to 1969. Psychol Med 31 (6):1045–1053.

Hovatta, I., J.D. Terwilliger, D. Lichtermann, T. Makikyro, J. Suvisaari et al., 1997 Schizophrenia in the genetic isolate of Finland. Am J Med Genet 74 (4):353–360.

Jakkula, E., K. Rehnstrom, T. Varilo, O.P. Pietilainen, T. Paunio et al., 2008 The genome-wide patterns of variation expose significant substructure in a founder population. Am J Hum Genet 83(6):787–794.

Jussila, A., L.J. Virta, V. Salomaa, J. Maki, A. Jula, and M. A. Färkkilä, 2013 High and increasing prevalence of inflammatory bowel disease in Finland with a clear North-South difference, J Crohns Colitis 7 (7):e256–262.

Kaipiainen-Seppanen, O., K. Aho, and M. Nikkarinen, 2001 Regional differences in the incidence of rheumatoid arthritis in Finland in 1995. Ann Rheum Dis 60 (2): 128–132.

Kerminen, S., A.S. Havulinna, G. Hellenthal, A.R. Martin, A.P. Sarin et al., 2017 Fine-Scale Genetic Structure in Finland. G3: Genes Genomes Genetics 7 (10):3459–3468.

Khera, A.V., M. Chaffin, K.G. Aragam, M.E. Haas, C. Roselli et al., 2018 Genome-wide polygenic scores for common diseases identify individuals with risk equivalent to monogenic mutations. Nat Genet 50 (9):1219–1224.

Kurki, M.I., E. Saarentaus, O. Pietilainen, P. Gormley, D. Lal et al., 2018 Contribution of rare and common variants to intellectual disability in a high-risk population sub-isolate of Northern Finland, bioRxiv, https://doi.org/10.1101/332023

Lango Allen, H., K. Estrada, G. Lettre, S.I. Berndt, M.N. Weedon et al., 2010 Hundreds of variants clustered in genomic loci and biological pathways affect human height. Nature 467 (7317):832–838.

Lappalainen, T., S. Koivumaki, E. Salmela, K. Huoponen, P. Sistonen et al., 2006 Regional differences among the finns: A Y-chromosomal perspective. Gene 376 (2):207–215.

Lawson, D.J., G. Hellenthal, S. Myers, and D. Falush, 2012 Inference of population structure using dense haplotype data. PLoS Genet 8 (1):e1002453.

Lehtinen, V., M. Joukamaa, K. Lahtela, R. Raitasalo, E. Jyrkinen et al., 1990 Prevalence of mental disorders among adults in Finland: basic results from the Mini Finland Health Survey. Acta Psychiatrica Scandinavica 81 (5):418–425.

Lehtinen, P., K. Pasanen, K. L. Kolho, and A. Auvinen, 2016 Incidence of Pediatric Inflammatory Bowel Disease in Finland: An Environmental Study. JPediatr GastroenterolNutr 63 (1):65–70.

Liu, J.Z., S. van Sommeren, H. Huang, S.C. Ng, R. Alberts et al., 2015 Association analyses identify 38 susceptibility loci for inflammatory bowel disease and highlight shared genetic risk across populations. Nat Genet 47 (9):979–986.

Locke, A.E., B. Kahali, S.I. Berndt, A.E. Justice, T.H. Pers et al., 2015 Genetic studies of body mass index yield new insights for obesity biology. Nature 518 (7538):197–206.

Loh, P.R., G. Tucker, B.K. Bulik-Sullivan, B.J. Vilhjalmsson, H.K. Finucane et al., 2015 Efficient Bayesian mixed-model analysis increases association power in large cohorts. Nat Genet 47 (3):284–290.

Marchini, J., L.R. Cardon, M.S. Phillips, and P. Donnelly, 2004 The effects of human population structure on large genetic association studies. Nat Genet 36 (5):512–517.

Martin, A.R., C.R. Gignoux, R.K. Walters, G.L. Wojcik, B.M. Neale et al., 2017 Human Demographic History Impacts Genetic Risk Prediction across Diverse Populations. Am J Hum Genet 100 (4):635–649.

Martin, A.R., M. Kanai, Y. Kamatani, Y. Okada, B.M. Neale et al., 2018a Hidden ‘risk’ in polygenic scores: clinical use today could exacerbate health disparities. bioRxiv, https://doi.org/10.1101/441261

Martin, A.R., K.J. Karczewski, S. Kerminen, M.I. Kurki, A.P. Sarin et al., 2018b Haplotype Sharing Provides Insights into Fine-Scale Population History and Disease in Finland. Am J Hum Genet 102 (5):760–775.

Mavaddat, N., P.D. Pharoah, K. Michailidou, J. Tyrer, M.N. Brook et al., 2015 Prediction of breast cancer risk based on profiling with common genetic variants. J Natl Cancer Inst 107 (5).

McEvoy, B.P., and P.M. Visscher, 2009 Genetics of human height. Econ Hum Biol 7 (3):294–306.

Neuvonen, A.M., M. Putkonen, S. Oversti, T. Sundell, P. Onkamo et al., 2015 Vestiges of an Ancient Border in the Contemporary Genetic Diversity of North-Eastern Europe. PLoS One 10(7):e0130331.

Nikpay, M., A. Goel, H.H. Won, L.M. Hall, C. Willenborg et al., 2015 A comprehensive 1,000 Genomes-based genome-wide association meta-analysis of coronary artery disease. Nat Genet 47 (10): 1121–1130.

O’Connell, K.S., N.W. McGregor, C. Lochner, R. Emsley, and L. Warnich, 2018 The genetic architecture of schizophrenia, bipolar disorder, obsessive-compulsive disorder and autism spectrum disorder. Mol Cell Neurosci 88:300–307.

Okada, Y., D. Wu, G. Trynka, T. Raj, C. Terao et al., 2014 Genetics of rheumatoid arthritis contributes to biology and drug discovery. Nature 506, 376–381.

Perala, J., S.I. Saarni, A. Ostamo, S. Pirkola, J. Haukka et al., 2008 Geographic variation and sociodemographic characteristics of psychotic disorders in Finland. Schizophr Res 106 (2-3):337–347.

Pietiläinen, O., 2014 Rare genomic deletions underlying schizophrenia and related neurodevelopmental disorders in Institute for Molecular Medicine Finland (FIMM) University of Helsinki, Helsinki.

Price, A.L., M.E. Weale, N. Patterson, S.R. Myers, A.C. Need et al., 2008 Long-range LD can confound genome scans in admixed populations. Am J Hum Genet 83 (1): 132–135; author reply 135–139.

Purcell, S., and C. Chang, PLINK 1.9. www.cog-genomics.org/plink/1.9/

Puska, P., E. Vartiainen, T. Laatikainen, P. Jousilahti, and M. Paavola, 2009 The Norther Karelia Project: From North Karelia To National Action. Helsinki: National Institute for Health and Welfare (THL), in collaboration with the North Karelia Project Foundation.

Reisberg, S., T. Iljasenko, K. Lall, K. Fischer, and J. Vilo, 2017 Comparing distributions of polygenic risk scores of type 2 diabetes and coronary heart disease within different populations. PLoS One 12 (7):e0179238.

Ripatti, P., J.T. Ramo, S. Soderlund, I. Surakka, N. Matikainen et al., 2016 The Contribution of GWAS Loci in Familial Dyslipidemias. PLoS Genet 12 (5):e1006078.

Salmela, E., T. Lappalainen, I. Fransson, P.M. Andersen, K. Dahlman-Wright et al., 2008 Genome-wide analysis of single nucleotide polymorphisms uncovers population structure in Northern Europe. PLoS One 3 (10):e3519.

Schizophrenia Working Group of the Psychiatric Genomics, C., 2014 Biological insights from 108 schizophrenia-associated genetic loci. Nature 511 (7510):421–427.

Schumacher, F.R., A.A. Al Olama, S.I. Berndt, S. Benlloch, M. Ahmed et al., 2018 Association analyses of more than 140,000 men identify 63 new prostate cancer susceptibility loci. Nat Genet 50 (7):928–936.

Scutari, M., I. Mackay, and D. Balding, 2016 Using Genetic Distance to Infer the Accuracy of Genomic Prediction. PLoS Genet 12 (9):e1006288.

Shungin, D., T.W. Winkler, D.C. Croteau-Chonka, T. Ferreira, A.E. Locke et al., 2015 New genetic loci link adipose and insulin biology to body fat distribution. Nature 518 (7538):187–196.

Silventoinen, K., S. Sammalisto, M. Perola, D.I. Boomsma, B.K. Cornes et al., 2003 Heritability of adult body height: a comparative study of twin cohorts in eight countries. Twin Res 6 (5):399–408.

Sohail, M., R.M. Maier, A. Ganna, A. Bloemendal, A.R. Martin et al., 2018 Signals of polygenic adaptation on height have been overestimated due to uncorrected population structure in genome-wide association studies. bioRxiv, https://doi.org/10.1101/355057

Stocker, H., T. Mollers, L. Perna, and H. Brenner, 2018 The genetic risk of Alzheimer’s disease beyond APOE epsilon4: systematic review of Alzheimer’s genetic risk scores. Transl Psychiatry 8 (1): 166.

R Core Team, 2018 R: A language and environment for statistical computing. R Foundation for Statistical Computing, Vienna, Austria. https://www.R-project.org/

THL, Sepelvaltimotauti-indeksi: Ikävakioitu (2013-2015). http://www.terveytemme.fi/sairastavuusindeksi/2015/maakunnat_html/atlas.html?select=01&indicator=i0

Visscher, P.M., N.R. Wray, Q. Zhang, P. Sklar, M.I. McCarthy et al., 2017 10 Years of GWAS Discovery: Biology, Function, and Translation. Am J Hum Genet 101 (1):5–22.

Wood, A.R., T. Esko, J. Yang, S. Vedantam, T.H. Pers et al., 2014 Defining the role of common variation in the genomic and biological architecture of adult human height. Nat Genet 46 (11): 1173–1186.

Yang, J., S.H. Lee, M.E. Goddard, and P.M. Visscher, 2011 GCTA: a tool for genome-wide complex trait analysis. Am J Hum Genet 88 (1):76–82.

Yengo, L., J. Yang, and P.M. Visscher, 2018 Expectation of the intercept from bivariate LD score regression in the presence of population stratification. bioRxiv, https://doi.org/10.1101/310565

