## Supplementary material for "Geographic variation and bias in polygenic scores of complex diseases and traits in Finland"

### Supplementary Information

### Supplemental Tables

**Table S1**. Results from the standard linear model without accounting for genetic relatedness. SNPs=number of variants in polygenic score (PS). East-West difference in PS is given in standard deviation unit of PS.

| | | Latitude | | Longitude | | |
|---|---|---|---|---|---|---|
| | **SNPs** | **Estimate** | **P-val** | **Estimate** | **P-val** | **E-W Difference (95% CI)** |
| **CAD** | 19,597 | 0.071 | 2.0e-6 | 0.11 | 4.2e-40 | 0.63 (0.55, 0,71) |
| **RA** | 32,736 | 0.10 | 4.4e-11 | 0.11 | 2.2e-41 | 0.63 (0.55, 0.63) |
| **CD** | 21,771 | 7.7e-3 | 6.2e-1 | -0.01 | 1.9e-1 | -0.10 (-0.19, -0.01) |
| **UC** | 23,513 | 0.07 | 2.6e-5 | 0.04 | 2.6e-7 | 0.26 (0.18, 0.35) |
| **SCZ** | 30,311 | 0.06 | 3.1e-4 | 0.07 | 3.2e-17 | 0.35 (0.26, 0.43) |
| **BMI** | 12,742 | 0.09 | 2.2e-8 | 0.09 | 8.1e-31 | 0.53 (0.44, 0.61) |
| **WHR** | 13,727 | 0.24 | 3.5e-60 | 0.18 | 4.0e-125 | 1.16 (1.09, 1.23) |
| **HG** | 27,066 | -0.35 | 2.7e-135 | -0.27 | 2.8e-320 | -1.51 (-1.58, -1.45) |

**Table S2.** Correlation with PS and PC1 and height variance explained ($R^2$) by PS before and after adjustment for PC1.

| | | $R^2$ | |
|---|---|---|---|
| | **Correlation with PC1** | **Before PC1 adjustment** | **After PC1 adjustment** |
| **GIANT-PS** | -0.798 | 0.1402 | 0.1916 |
| **UKBB-PS** | -0.103 | 0.2229 | 0.2118 |
| **FINRISK-PS** | -0.304 | 0.1542 | 0.1352 |

**Table S3.** Comparison of PS for HG using GWAS results based on standard linear model adjusted for 10 principal components or linear mixed model in FINRISK and UK Biobank data.

| | FINRISK | | UK Biobank | |
|---|---|---|---|---|
| | **$R^2$** | **WF-EF difference (95% CI, cm)** | **$R^2$** | **WF-EF difference (95% CI, cm)** |
| **Standard linear model** | 15% | 1.35 (1.14 – 1.58) | 22% | 0.64 (0.39 – 0.89) |
| **Linear mixed model** | 15% | 1.15 (0.94 – 1.38) | 25% | 0.37 (0.10 - 0.64) |

**Table S4.** Comparison of Finns-excluded and Finns-included GWAS in schizophrenia. Results are shown for the linear model similarly to Table 1 in the main text. SNPs=number of variants in polygenic score (PS). Difference in PS between EF and WF subpopulations is given in standard deviation unit of PS.

| | | **Latitude** | | **Longitude** | | |
|---|---|---|---|---|---|---|
| | **SNPs** | **Estimate** | **P-val** | **Estimate** | **P-val** | **EF-WF Difference (95% CI)** |
| **Finns excluded** | 30,311 | 0.04 | 8.7e-2 | 0.04 | 4.0e-3* | 0.35 (0.26, 0.43) |
| **Finns included** | 30,760 | 0.10 | 3.1e-6* | 0.05 | 2.6e-4* | 0.41 (0.32, 0.49) |

**Table S5.** Linear regression results for explaining height with different PS in eastern and western subpopulations separately.

| | **$R^2$** | | | **E-W difference (cm)** | | |
|---|---|---|---|---|---|---|
| | **All (n=2,373)** | **East (n=1601)** | **West (n=772)** | **All** | **East** | **West** |
| **GIANT-PS** | 0.1402 | 0.1771 | 0.1727 | -3.5 (-3.9, -3.1) | -6.4 (-7.1, -5.7) | -4.7 (-5.4, -3.9) |
| **UKBB-PS** | 0.2229 | 0.2126 | 0.2241 | -0.6 (-0.9, -0.4) | -0.6 (-0.9, -0.4) | -0.6 (-0.9, -0.4) |
| **FINRISK-PS** | 0.1542 | 0.1581 | 0.104 | -1.4 (-1.6, -1.1) | -1.3 (-1.6, -1.1) | -1.3 (-1.6, -1.0) |

**Table S6.** Comparison of BMI PS. (Similar to Table 1 and Table S2 for HG.)

| | | | **Latitude** | | **Longitude** | | |
|---|---|---|---|---|---|---|---|
| | **SNPs** | **$R^2$** | **Estimate** | **P-val** | **Estimate** | **P-val** | **E-W Difference (95% CI)** |
| **GIANT-PS** | 12,742 | 8.0 % | 0.03 | 0.094 | 0.04 | 1.8e-3 | 0.64 (0.51, 0.77) |
| **UKBB-PS** | 75,979 | 4.9 % | 0.01 | 0.60 | 0.01 | 0.67 | 0.05 (-0.03, 0.14) |
| **FINRISK-PS** | 44,920 | 1.3 % | -0.01 | 0.80 | -0.002 | 0.90 | 0.08 (0.04, 0.14) |

| | | **$R^2$** | |
|---|---|---|---|
| | **Correlation with PC1** | **Before PC1 adjustment** | **After PC1 adjustment** |
| **GIANT-PS** | 0.286 | 8.0 % | 7.8 % |
| **UKBB-PS** | 0.048 | 4.9 % | 4.8 % |
| **FINRISK-PS** | 0.098 | 1.3 % | 1.2 % |

**Table S7.** Comparison of WHR PS. (Similar to Table 1 and Table S2 for HG.)

| | | | Latitude | | Longitude | | |
|---|---|---|---|---|---|---|---|
| | **SNPs** | **$R^2$** | **Estimate** | **P-val** | **Estimate** | **P-val** | **E-W Difference (95% CI)** |
| **GIANT-PS** | 13,130 | 2.0 % | 0.10 | 1.0e-9 | 0.08 | 4.7e-12 | 0.007 (0.005, 0.01) |
| **FINRISK-PS** | 43,252 | 1.1 % | -0.11 | 0.63 | -0.03 | 0.017 | -5e-4 (-1e-3, -1e-4) |

| | | $R^2$ | |
|---|---|---|---|
| | **Correlation with PC1** | **Before PC1 adjustment** | **After PC1 adjustment** |
| **GIANT-PS** | 0.58 | 2.0 % | 2.2 % |
| **FINRISK-PS** | -0.04 | 1.1 % | 1.1 % |

#### Supplemental Figures

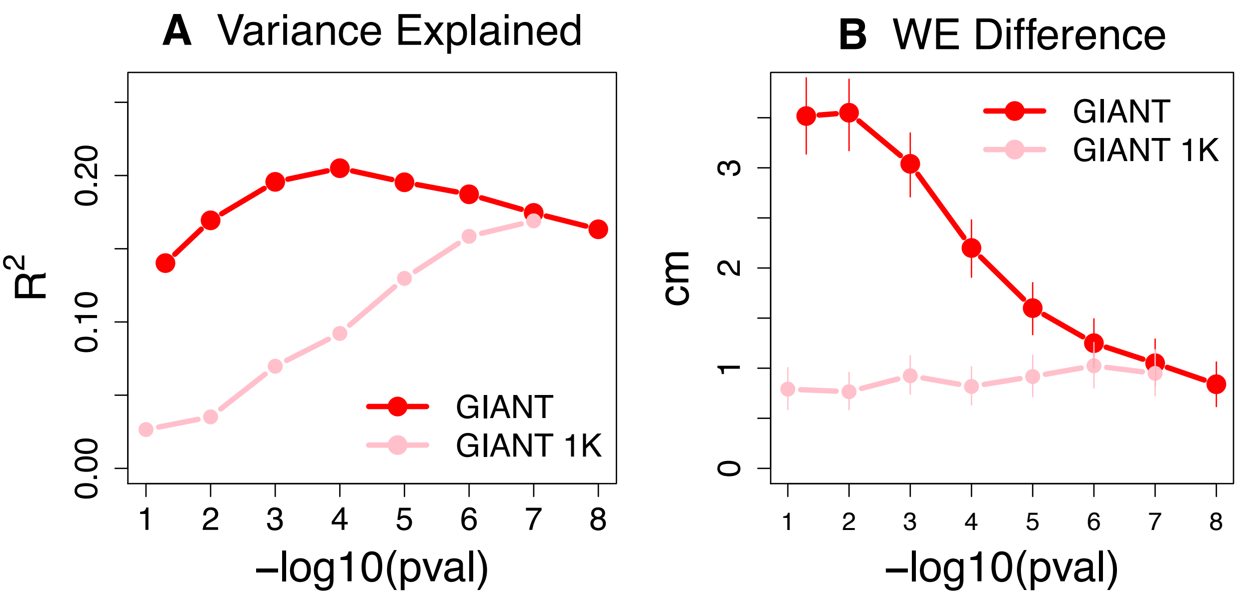

**Figure S1. A)** Height variance explained by GIANT-PS and a subset of GIANT-PS with at most 1,000 randomly sampled variants, as a function of P-value threshold in GIANT data. **B)** predicted West-East difference in height by the two PS, as a function of P-value threshold in GIANT data. Variance explained is given as adjusted $R^2$.

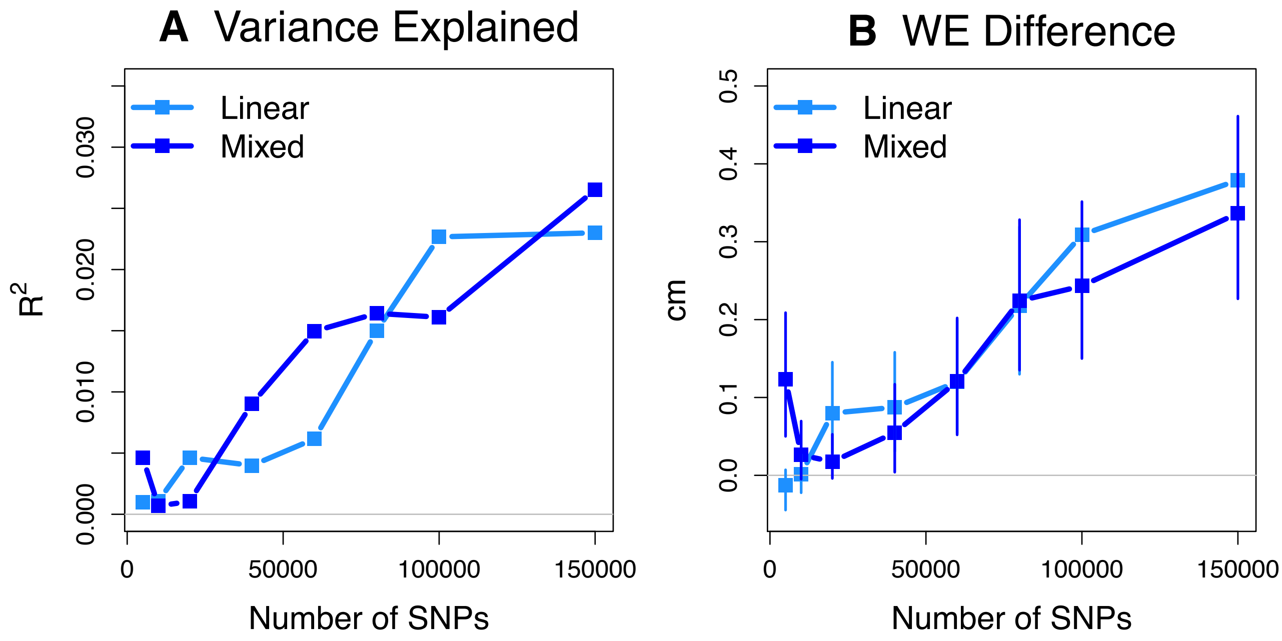

**Figure S2.** A comparison between random PS (i.e. all variants have P-value > 0.5 in GWAS) from the height GWAS on the FINRISK cohort ran with standard linear model or linear mixed model. **A)** Height variance explained by PS and **B)** predicted West-East difference in height by PS, as a function of the number of independent variants in PS. Variance explained is given as adjusted $R^2$.

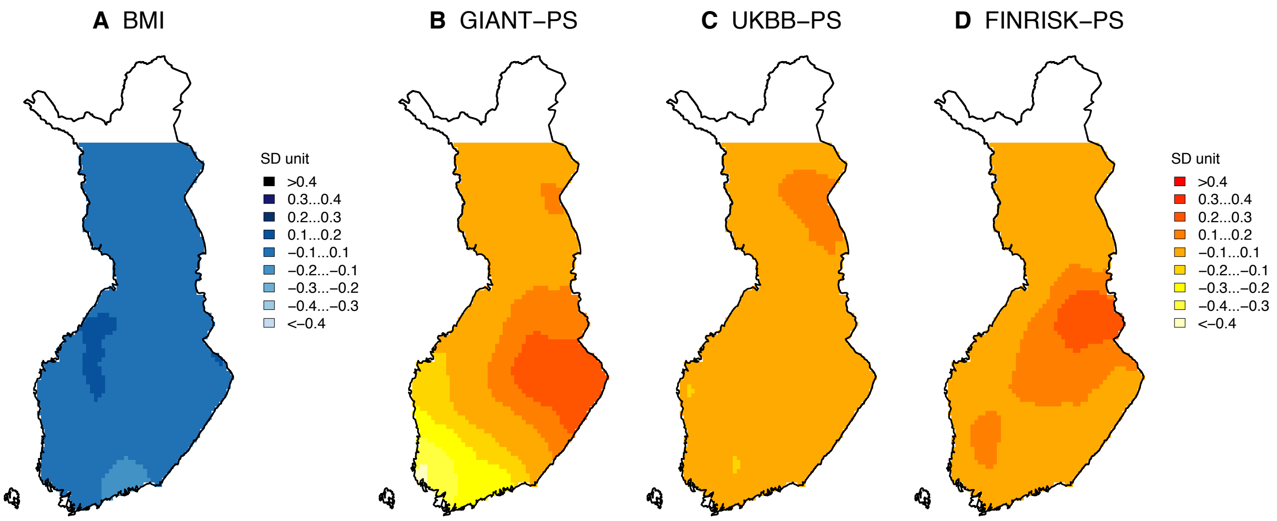

**Figure S3. A)** Distribution of sex, age and age² adjusted BMI and polygenic score (PS) distributions of **B)** GIANT-PS, **C)** UKBB-PS and **D)** FINRISK-PS for BMI in Finland. The values are in standard deviation unit. The observed BMI does not show differences between subpopulations (95% CI: -0.12, 0.59).

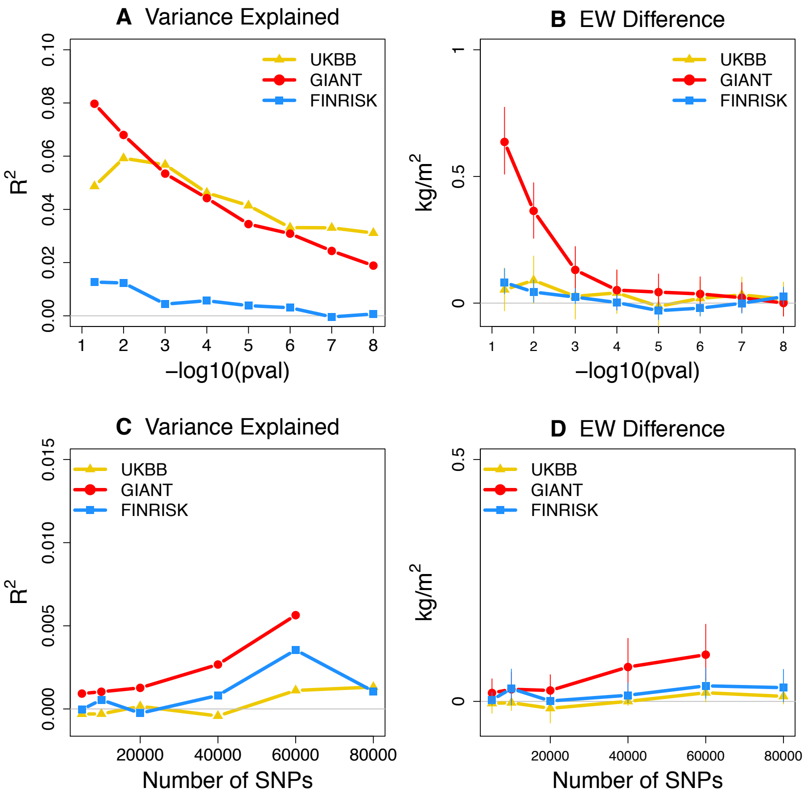

**Figure S4**. Top row: **A)** BMI variance explained by PS and **B)** predicted East-West difference in BMI by PS, as a function of P-value threshold in GWAS data.
Bottom row: **C)** BMI variance explained by PS and **D)** predicted East-West difference in BMI by PS, as a function of the number of independent variants in PS when all variants have P-value > 0.5 in GWAS. Variance explained is given as adjusted $R^2$.

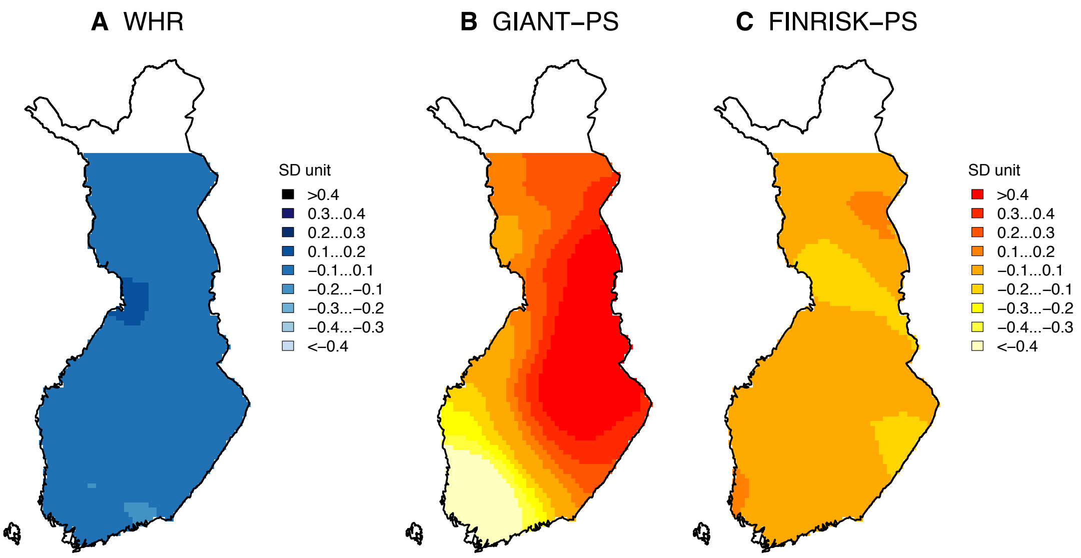

**Figure S5. A)** Distribution of sex, age, age[2] and BMI adjusted WHR and polygenic score (PS) distributions of **B)** GIANT-PS, **C)** FINRISK-PS for WHR (adjusted for BMI) in Finland. The values are in standard deviation unit. The observed WHR (adjusted for BMI) does not show differences between subpopulations (95% CI: −0.0009, 0.0072).

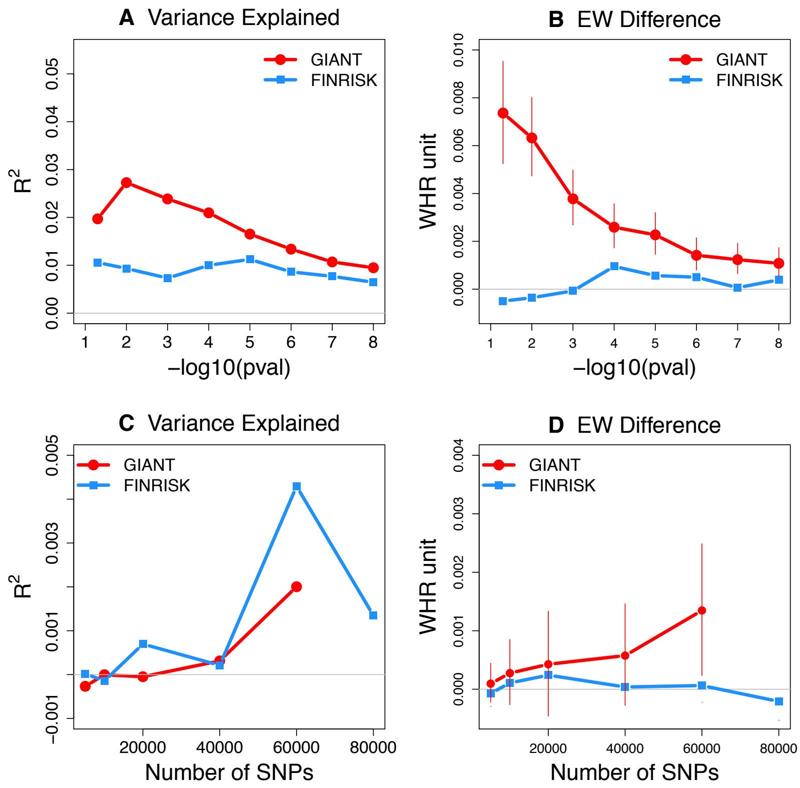

**Figure S6**. Top row: **A)** WHR variance explained by PS and **B)** predicted East-West difference in WHR (adjusted for BMI) by PS, as a function of P-value threshold in GWAS data.
Bottom row: **C)** WHR variance explained by PS and **D)** predicted East-West difference in WHR (adjusted for BMI) by PS, as a function of the number of independent variants in PS when all variants have P-value > 0.5 in GWAS. 95% CI is so small for FINRISK that it is not visible in the figure. Variance explained is given as adjusted $R^2$.

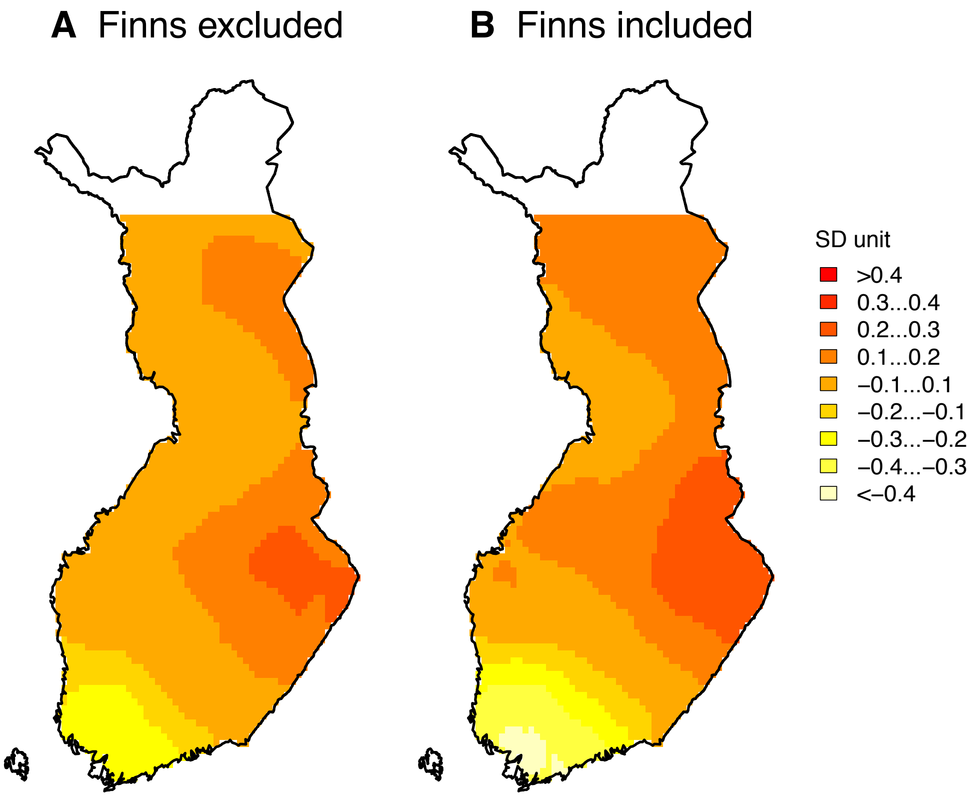

**Figure S7.** Distribution of polygenic scores for schizophrenia based on GWAS **A)** excluding Finnish samples and **B)** including Finnish samples.

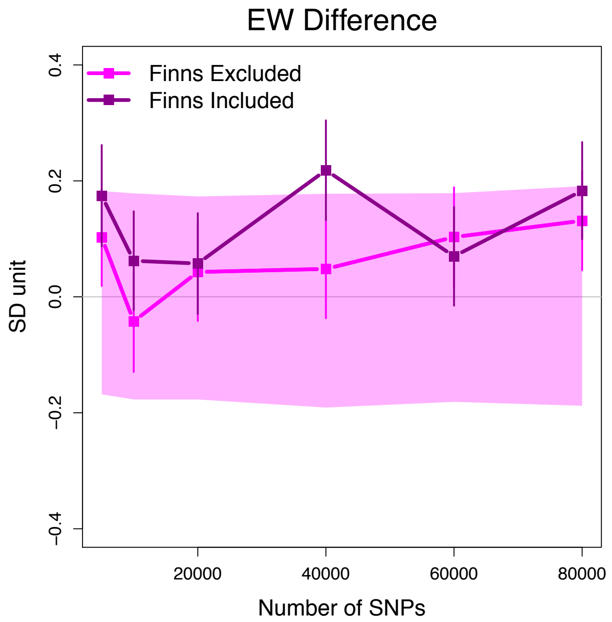

**Figure S8.** PS differences between Eastern and Western subpopulations using different numbers of independent variants randomly chosen with GWAS P-value > 0.5. Results for PS built based on SCZ GWAS with excluding (magenta) and including (dark violet) Finnish samples do not show major accumulation of geographic differences.

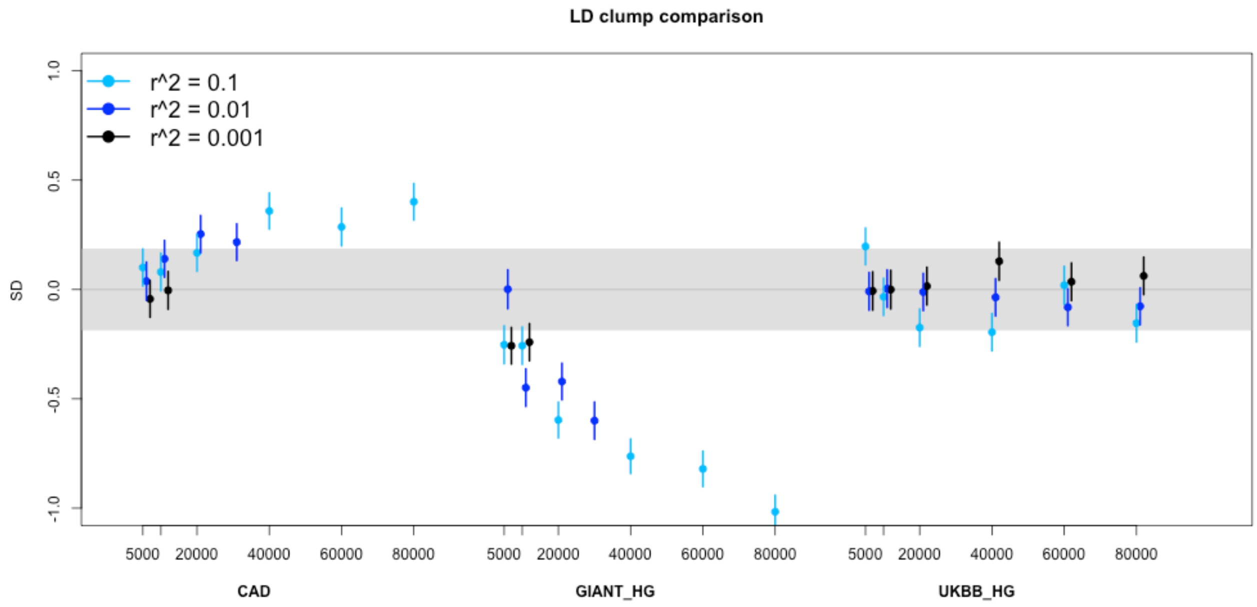

**Figure S9.** Comparison of different $r^2$ thresholds on differences between Eastern and Western subpopulations in randomly chosen variants with GWAS P-value > 0.5. The solid region is the 95% probability interval under the theoretical null assumption of zero effect sizes and completely independent variants ($r^2 = 0$).

#### SUPPLEMENTAL TEXT : DISTRIBUTION OF POLYGENIC SCORE DIFFERENCE

Consider $M$ independent variants and assume that standard error of effect size estimate of variant $k$ is $s_k$. For both quantitative traits and diseases, $s_k \approx c \cdot (2f_k(1-f_k))^{-\frac{1}{2}}$, where $f_k$ is the minor allele frequency of variant $k$ and $c$ is a constant that depends on the sample size and observed phenotypic variance in the GWAS data but is independent of $k$.

Assume that true effects are zero at these variants whence $\widehat{\beta}_k \sim \mathcal{N}(0, s_k^2)$. Denote the z-score by $\widehat{z}_k = \widehat{\beta}_k / s_k \sim \mathcal{N}(0,1)$.

The polygenic score for individual $i$ is

$$p_i = \sum_{k=1}^{M} g_{ik} \widehat{\beta}_k = \sum_{k=1}^{M} g_{ik} s_k \widehat{z}_k,$$

where $g_{ik}$ is the effect allele dosage of $i$ at variant $k$. By mean-centering the score, we may assume that the allele dosages were mean-centered at every variant.

Since effect estimates and genotypes are independent, $\mathrm{E}(p_i) = 0$ and

$$\mathrm{Var}(p_i) = \sum_{k=1}^{M} \mathrm{E}(g_{ik}^2 s_k^2 \widehat{z}_k^2) = \sum_{k=1}^{M} 2f_k(1-f_k) \cdot s_k^2 \cdot 1 = Mc^2.$$

Hence, the standardized score is $p_i^* = p_i / (\sqrt{M}c)$.

Consider the difference $\Delta$ between scores of a Western and an Eastern individual:

$$\Delta = p_W^* - p_E^* = \sum_{k=1}^{M} \frac{(g_{Wk} - g_{Ek})}{\sqrt{M}c} s_k \widehat{z}_k = \sum_{k=1}^{M} \frac{(g_{Wk} - g_{Ek})}{\sqrt{2Mf_k(1-f_k)}} \widehat{z}_k.$$

$\mathrm{E}(\Delta) = 0$ because $\mathrm{E}(\widehat{z}_k) = 0$ and

$$\mathrm{Var}(\Delta) = \sum_{k=1}^{M} \frac{\mathrm{Var}((g_{Wk} - g_{Ek})\widehat{z}_k)}{2Mf_k(1-f_k)} = \mathrm{Var}\left( \frac{(g_{Wk} - g_{Ek})\widehat{z}_k}{\sqrt{2f_k(1-f_k)}} \right).$$

Variance of $\Delta$ depends only on the second moment of the distribution of standardized dosage difference between West and East. In particular, variance of $\Delta$ is independent of $M$ and sample size and phenotypic variance of the GWAS from which the effect size estimates come from. This explains why the 95% regions of $\Delta$ appear approximately constant across all traits and diseases considered.
